# Episodic memory in aspects of brain information transfer by resting-state network topology

**DOI:** 10.1101/2021.02.28.433300

**Authors:** Tianyi Yan, Gongshu Wang, Li Wang, Tiantian Liu, Ting Li, Luyao Wang, Dingjie Suo, Shintaro Funahashi, Duanduan Chen, Bin Wang, Jinglong Wu

**Affiliations:** School of Life Science, Beijing Institute of Technology, Beijing, China, 100081; Intelligent Robotics Institute, School of Mechatronical Engineering, Beijing Institute of Technology, Beijing 100081; Advanced Research Institute of Multidisciplinary Science, Beijing Institute of Technology, Beijing, China 100081; Department of Information and Computer, Taiyuan University of Technology Taiyuan, Shanxi 030024; Key Laboratory of Biomimetic Robots and Systems, Ministry of Education, Beijing, China, 100081; International Joint Research Laboratory of Biomimetic Robots and Systems, Ministry of Education, Beijing, China, 100081

## Abstract

Studies suggest that resting-state functional connectivity conveys cognitive information; also, activity flow mediates cognitive information transfer. However, the exact mechanism of interregional interactions underlying episodic memory remains unclear. We performed a combined analysis of task-evoked activity and resting-state functional connectivity by activity flow mapping to estimate the information transfer mechanism of episodic memory. We found that the cognitive control and attentional networks were the most recruited structures in information transfers during both encoding and retrieval processes; these networks were correlated with task-evoked activation. Differences in information transfer intensity between encoding and retrieval mainly existed in the visual, somatomotor and hippocampal systems. Furthermore, information transfer showed high predictive power for episodic memory ability and mediated relationships between task-evoked activation and memory performance. Additional analysis indicated that structural connectivity had a transportive role in information transfer. Finally, our study presented the information transfer mechanism of episodic memory from multiple neural perspectives.

## Introduction

Episodic memory is critical for human survival. Episodic memory includes the memory of personal experiences of life events (Tulving 2002, Rosenbaum, Stuss et al. 2007) and consists of encoding and retrieval processes. Encoding refers to the conversion of experiences into stored information, and retrieval refers to the recall of stored information. Task-based studies employing functional magnetic resonance imaging (fMRI) revealed that both encoding and retrieval induced activation in the medial temporal lobe (MTL) (Staresina and Davachi 2008, Kazumasa Z. Tanaka 2014), prefrontal cortical region (Shigemune, Tsukiura et al. 2017) and parietal cortical region (Cabeza, Ciaramelli et al. 2008, Hutchinson, Uncapher et al. 2014).

More recent studies suggest that episodic memory involves communication between discrete regions. Brain network analysis in healthy young and old adults (Woorim, Kee et al. 2015, Campbell, Madore et al. 2018) and in patients with memory impairments (Sidhu, Jason et al. 2013, Bryce, Mander et al. 2015) all indicated the involvement of interregional interactions in episodic memory (Friston and J. 2011, Park and Friston 2013). In a retrieval task, an interaction between the medial temporal lobe and the prefrontal and parietal cortex has been consistently reported (Geib, Stanley et al. 2015, Cooper, Richter et al. 2017). Furthermore, the functional coupling between temporal and frontoparietal regions was related to true recognition, whereas a more distributed hippocampal-temporal-parietal-prefrontal network was related to false recognition (Paz-Alonso, Gallego et al. 2013, Blankenship, Redcay et al. 2016). However, these studies established functional connectivity (FC) through the association between regional signals. The relatively ambiguous estimates of causal paths between regions fail to determine effective interactions that accurately encode cognitive information (Ito, Hearne et al. 2019, Reid, Headley et al. 2019).

Studies have suggested that resting-state FC elucidates the intrinsic network architecture and can characterize the individual performance of episodic memory (Vincent, Patel et al. 2007, Sala-Llonch, Pe?a-Gómez et al. 2016, Chén, Cao et al. 2019). The evidence indicates a positive association among the default mode network (DMN), and DMN-hippocampus connectivity and memory performance (Ward, Mormino et al. 2015, Kaboodvand, Bäckman et al. 2018). Moreover, positive correlations between resting-state FCs in the anterior temporal structure (i.e., the temporal pole, the amygdala, the entorhinal cortex, and the hippocampus) and memory performance have been observed across people of different ages (Grothe and Teipel 2016).

Studies have revealed significant associations between resting-state FC and task-evoked activation during encoding and retrieval tasks (Hermundstad, Bassett et al. 2013, Ritchey, Yonelinas et al. 2014). Recently, Cole et al. proposed a new approach of activity flow mapping by using resting-state FCs to explain task-evoked activity and designed a model to estimate information transfer during logic, sensory, and motor tasks. These results confirmed that resting-state FC allows for the transmission of cognitive information and that activity flow mediates decodable cognitive information transfer between brain regions. Therefore, a combined analysis of resting-state FC and task-evoked activation based on activity flow mapping could estimate cognitive-related information interactions between regions and encode brain functional networks more accurately, which is important for complex cognitive functions such as episodic memory (Cole, Ito et al. 2016, Ito, Kulkarni et al. 2017, Ito, Hearne et al. 2019).

Based on the above consideration, we combined resting-state FC with task-evoked activation to elucidate the brain network patterns of information transfer underlying episodic memory. First, we computed the information transfer intensity between each paired region and compared their differences during encoding and retrieval. Then, we explored the relationship between information transfer with task-evoked activation and memory performance and investigated whether there was a mediating effect among these factors. In addition, we analyzed the effect of structural connections (SCs) on information transfer and verified the results from multiple aspects of pathological memory, brain state changes and simulation models.

## Results

### Information transfer during encoding and retrieval

We first mapped the information transfer (see methods for details) between regions during an episodic memory task at the whole-brain level. As shown in Figure 2a, there are 4313 effective transfers in encoding and 4490 effective transfers in retrieval (for a total of 129240 possible transfers). Figure 2b shows the percent of statistically effective information transfers for each paired network (displaying the results of instances when the number of transfers was larger than 2; the number of effective transfers per pair of networks is shown in Figure 2—Supplementary figure 1). It can be seen that the distributions of these transfers were not random. During encoding, the region-to-region transfer was highly distributed involving all the networks, and most transfers were among and within the cingulo-opercular network (CON), DMN, and frontoparietal network (FPN); more specifically, the transfers were within the visual network (VIS), auditory network (AUD), dorsal attention network (DAN) and hippocampal system (HIPP) and were among the VIS, FPN and DAN. During retrieval, the region-to-region transfers were also highly distributed within the somatomotor network (SMN), CON, DMN, FPN, AUD and DAN and among the CON, DMN, FPN and DAN. After sorting the data by the number of transfers, we found that the CON, DMN, FPN and DAN demonstrated the most effective information transfer both in the internetwork and intranetwork during encoding (68.9% of all information) and retrieval (90.2% of all information).

**Figure 1.**
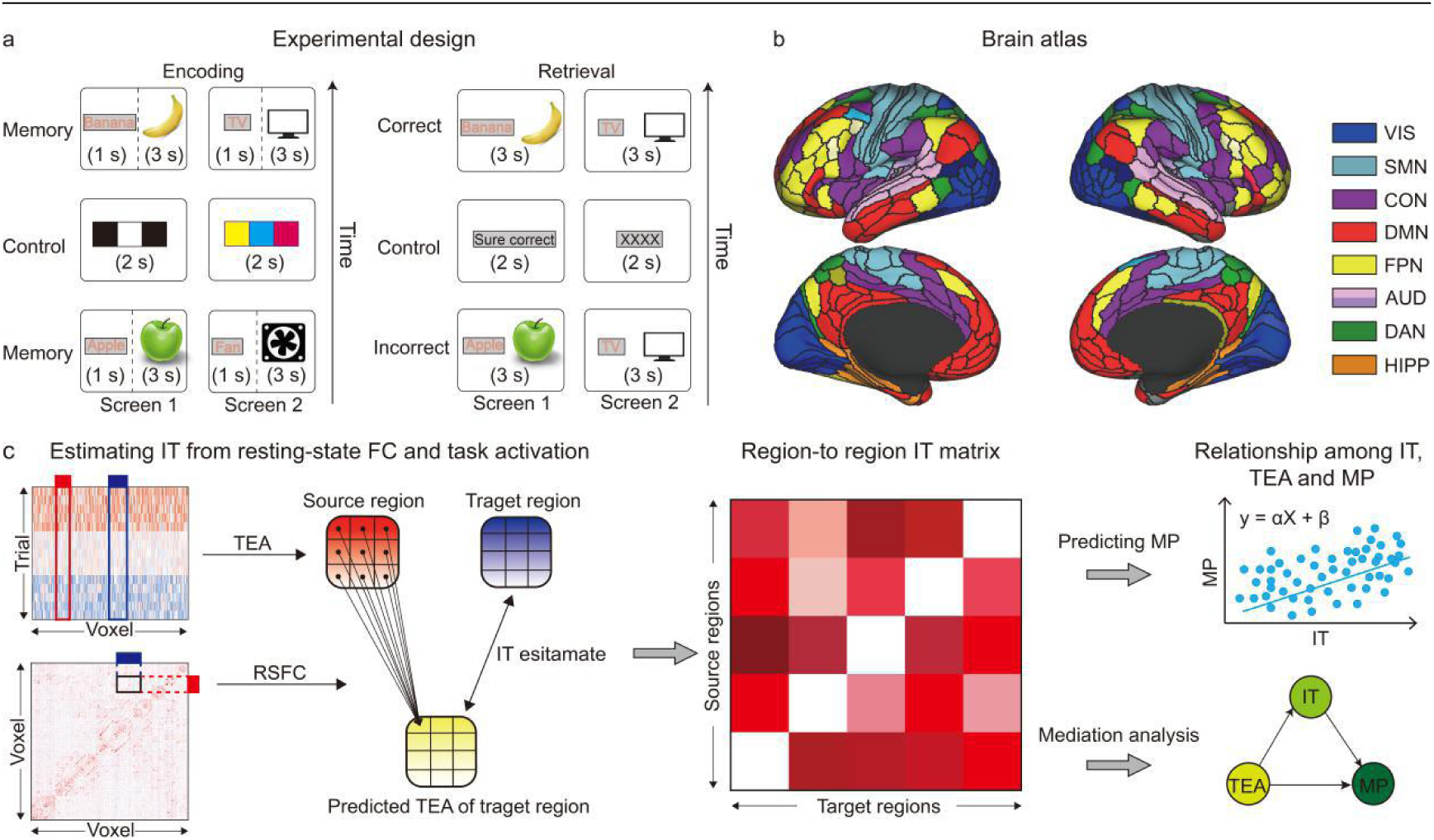
Task paradigm design and experimental process. **a** Example trial for each task. The encoding task included 40 memory trials and 24 control trials. In memory trials, a combination of a word and a matched image appeared on each screen. Participants were asked to remember the paired objects on the screens. In control trials, one screen appeared black or white, and the other appeared in color. Retrieval tasks included 40 correct trials, 40 incorrect trials and 24 control trials. In correct trials, items were shown to be paired as they had been at the encoding stage. In incorrect trials, items were shown in different pairs than they were in at the encoding stage. Subjects were required to look at a pair of objects and rate their confidence in their memory of the pairing. During control trials, one side of the screen displayed one of the four retrieval confidence response options, and the other side of the screen was “xxxx”. **b** Glass’s brain atlas was used to parcel the cerebral cortex into 360 regions and then assign each region to a functional subnetwork. This study focused on 8 main subnetworks. **c** The contents of this study. First, information transfer mapping was used to estimate the information transfer per pair of regions and to calculate the information transfer intensity of each region. Then, the correlation between regional information transfer intensity and task-evoked activation and that between regional information transfer intensity and memory performance were analyzed; then, the predictive power of the information transfer matrix on memory performance was texted. Finally, whether mediation was evident among task-evoked activation, information transfer intensity and memory performance was explored. (RSFC=resting-state FC, TEA=task-evoked activation, IT=information transfer, MP=memory performance)

**Figure 2.**
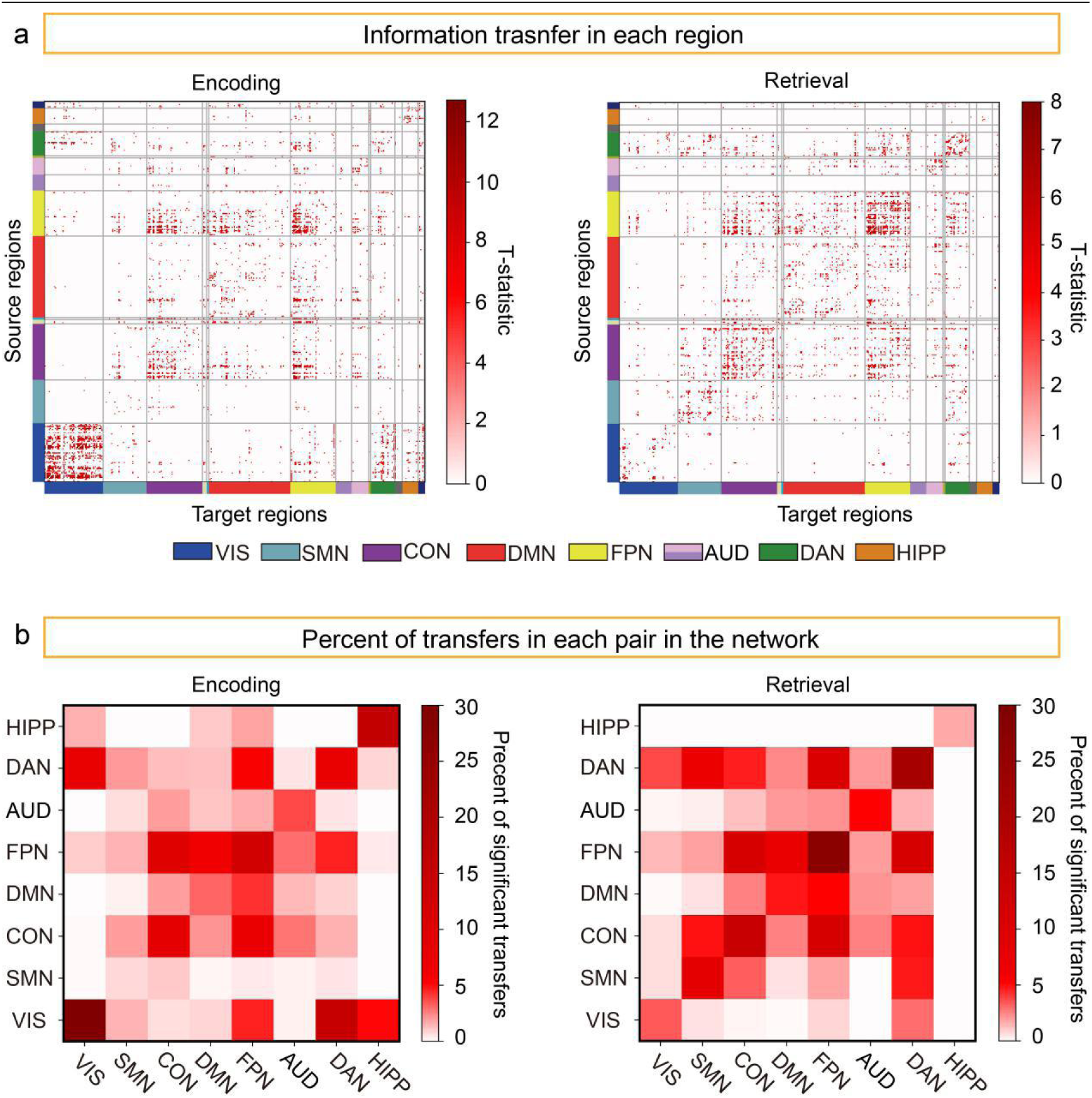
Information transfer estimation between all pairs of regions. All reported results were statistically significant at a threshold of p<0.05 (FDR correction). **a** Region-to-region information transfer mapping during encoding and retrieval. **b** Percentage of effective region-to-region information transfers by network affiliation in a and b to better visualize and assess how region-to-region transfer mappings may have been influenced by underlying network organization. Only the results in which the number of transfers of network affiliation was larger than 2 are displayed.

### Brain regions involving information transfers

We then calculated the percent of statistically effective information transfers of each cortical region to visualize the ability of regions to initiate information transfer. Figure 3 shows the results and displays only regions in which the percentage was larger than 0.15% (equal to a transfer number greater than 2), with 278 regions involved in encoding and 271 regions involved in retrieval (360 regions in total). The encoding and retrieval regions showed different but overlapping results. The areas with massive transfers (approximately the top 20%) were mainly concentrated in the paracentral lobule, the midcingulate cortex, the premotor cortex, and the dorsolateral prefrontal cortex of the CON and in the inferior parietal cortex, the anterior cingulate cortex, the medial prefrontal cortex, the inferior frontal cortex, and the dorsolateral prefrontal cortex of the FPN during both encoding and retrieval. Furthermore, we observed many distinctions between encoding and retrieval.

**Figure 3.**
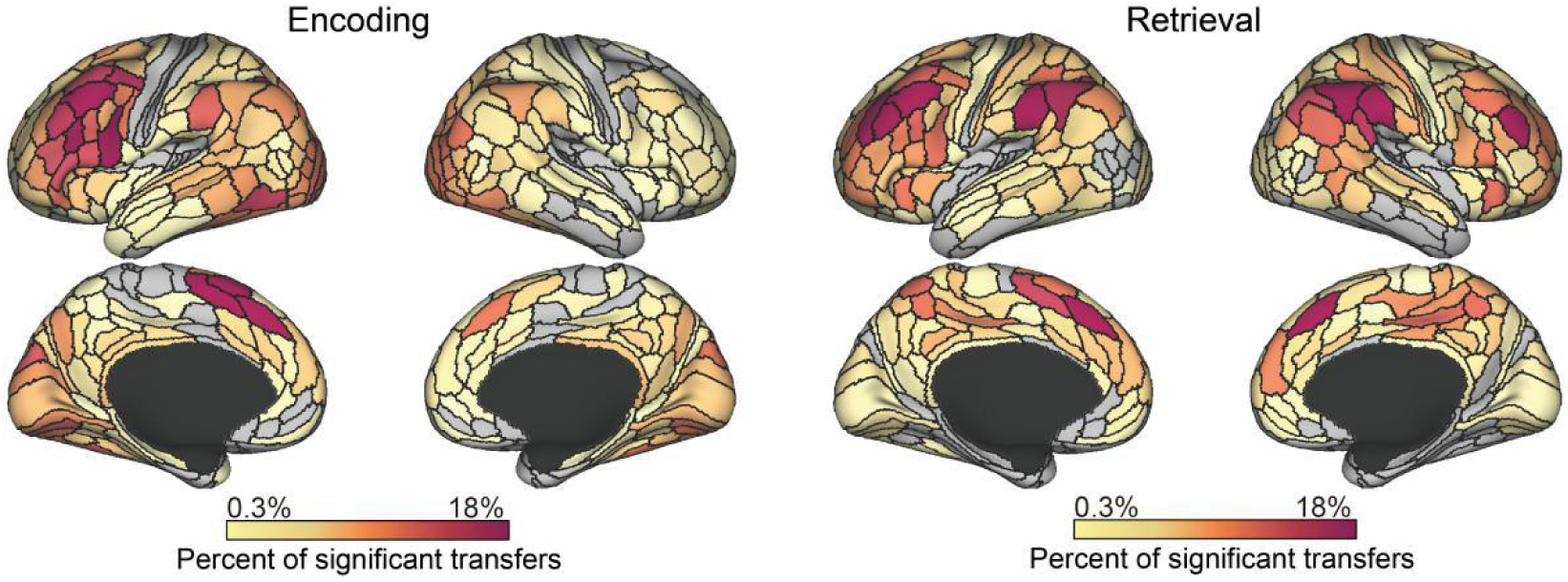
Percent of effective information transfers of each cortical region during encoding and retrieval. All reported information transfers were statistically significant at p < 0.05 with FDR correction. Percentages were computed by dividing the number of significant transfers from each region by the total number of possible transfers from that region (359×2 other regions). Only the regions where the transfer number was larger than 2 are displayed.

Except for the above areas, the active areas during encoding were mainly located in the whole VIS, the inferior parietal cortex of the DAN, and the parahippocampal gyrus of the HIPP, whereas the active areas during retrieval were mainly in the inferior parietal cortex and inferior frontal cortex of the CON; the insular and frontal opercular cortex of the FPN; the temporo-parieto-occipital junction of the AUD; and the lateral temporal and superior parietal cortex and posterior cingulate cortex of the DAN (Figure 3).

### Differences in information transfer between encoding and retrieval

Given that encoding and retrieval are two different processes in episodic memory, we compared them with regard to information transfer. As shown in Figure 4a, the information transfer of the VIS and HIPP was significantly stronger in encoding than in retrieval (t_(79)_ = 12.91, p = 2.1 × 10^-20^; t_(79)_ = 10.16, p = 3.8×10^-15^, respectively), whereas the SMN was significantly stronger during retrieval than during encoding (t_(79)_ = -5.90, p = 6.5×10^-7^). We further compared encoding and retrieval at a regional level and found that the information transfer intensity of 123 regions (VIS:52, SMN:3, CON:11, DMN:13, FPN:7, AUD:10, DAN:6, and HIPP:14) was stronger during encoding (yellow areas) and that of 102 regions (VIS:1, SMN:20, CON:17, DMN:20, FPN:20, AUD:8, DAN:11, and HIPP:0) was stronger during retrieval (blue areas) (p<0.05 with Bonferroni correction).

**Figure 4.**
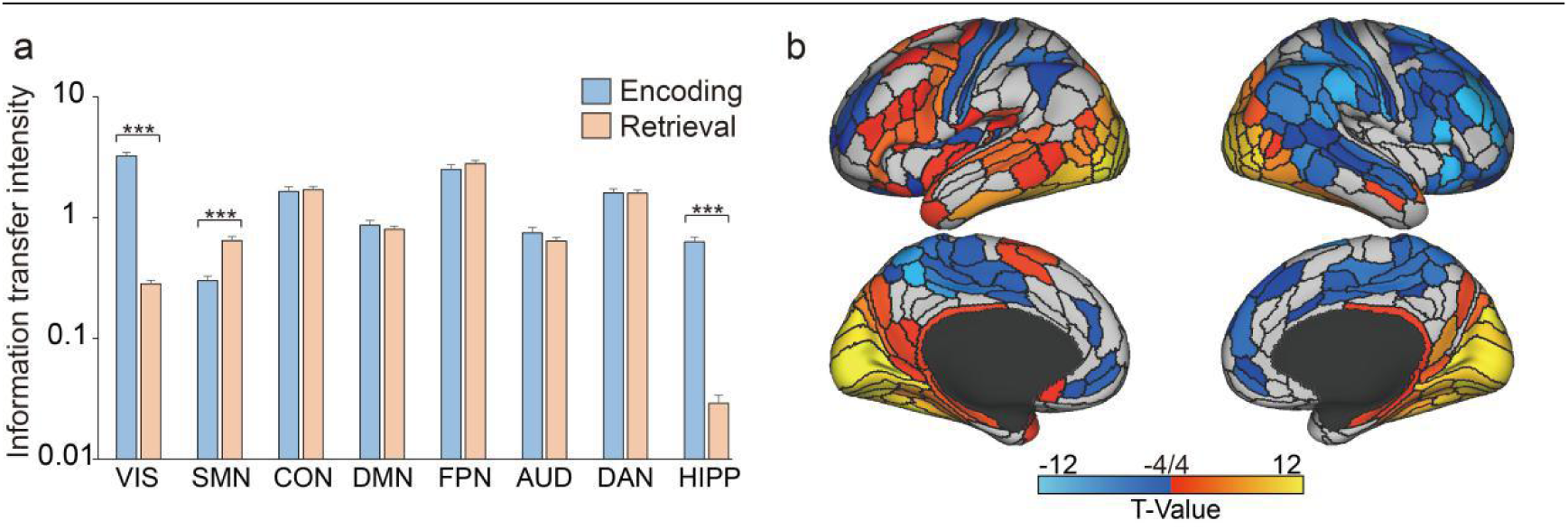
Difference in information transfer intensity between encoding and retrieval. The significance of the difference was measured by a threshold of p < 0.05 with Bonferroni correction. **a** Average information transfer intensity for each functional subnetwork. **b** Difference in the information transfer intensity of each region between encoding and retrieval. Blue indicates that information transfer intensity was stronger in retrieval than in encoding, and red indicates that the information transfer intensity was stronger in encoding than in retrieval. *** indicates p<0.001 with Bonferroni correction; error bars indicate the SE of estimates.

We observed that information transfer of regions in the primary visual cortex, early visual cortex, dorsal stream visual cortex, ventral stream visual cortex, MT+ complex and neighboring visual areas, medial temporal cortex, lateral temporal cortex, and posterior cingulate cortex were stronger during encoding; also, information transfer of regions in the somatosensory and motor cortex, paracentral lobular and midcingulate cortex, superior parietal cortex, anterior cingulate cortex, medial prefrontal cortex, and dorsolateral prefrontal cortex were stronger during retrieval (p<0.05 with Bonferroni correction, Figure 4b).

### Relationship between information transfer and task-evoked activation

Given that task-evoked activation represents the metabolism level and that information transfer intensity reflects the signal transmission ability, we then investigated whether there is an association between these two properties.

Figure 5 illustrates the regions where information transfer intensity was significantly correlated with task-evoked activation (Pearson’ |r| > 0.29, p < 0.01), with 107 regions involved in encoding and 67 regions involved in retrieval. We observed that the distribution of these regions was similar during both encoding and retrieval. During encoding, most of the positively correlated regions were located in the FPN and the CON (mainly in the anterior cingulate and inferior frontal cortices), whereas most negatively correlated regions were in the posterior cingulate cortex (the DMN) (Figure 5). During retrieval, the positively correlated regions were also located in the FPN and the CON, whereas most negatively correlated regions were in the posterior cingulate (CON) and inferior parietal cortex (DMN) (Figure 5). In addition, functional activity analysis results showed that almost all the negatively correlated regions were deactivated (Figure 5—Supplementary figure 1).

**Figure 5.**
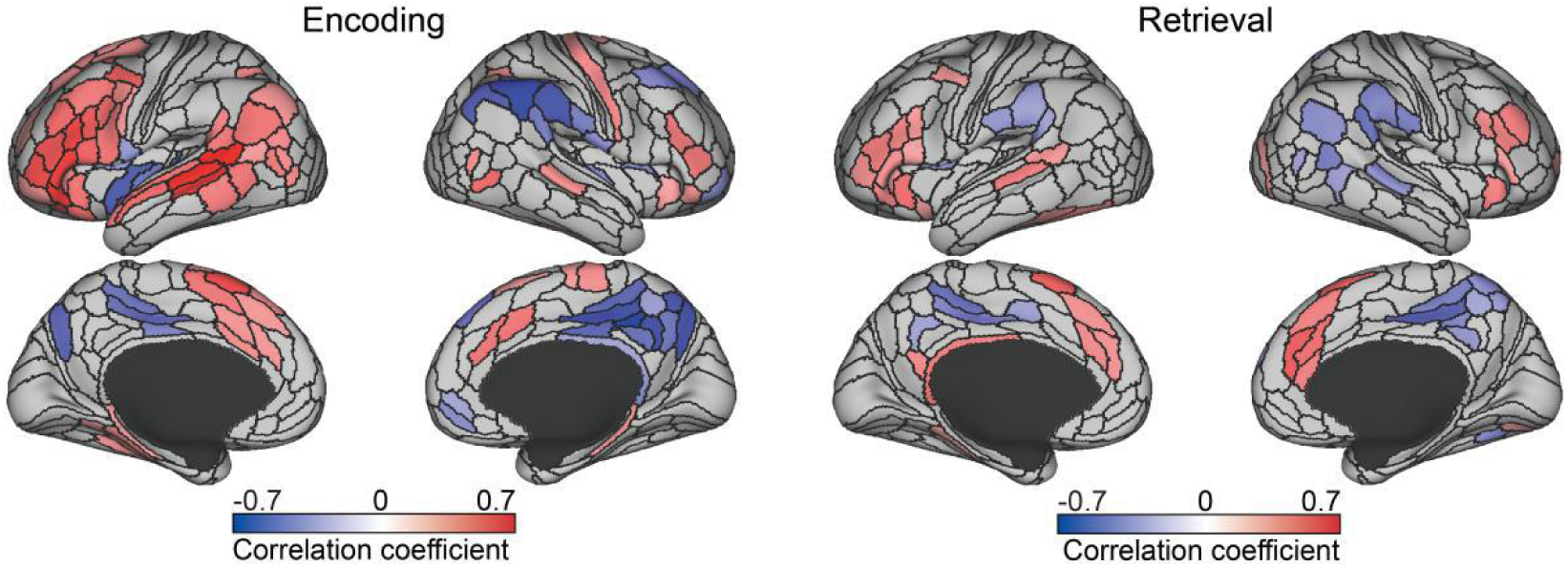
The correlation between information transfer intensity and averaged task-evoked activation in each region during encoding and retrieval. All reported regions were statistically significant at p < 0.01. Information transfer intensity was computed by the weighted sum of transfers from and to each region, and averaged task-evoked activation was computed by the average beta value of all trials for each region.

### The relationship between information transfer and memory ability

Communication between brain regions contributes to cognitive function. Then, we investigated whether the information transfer in episodic memory influences memory performance. As shown in Figure 6, information transfer intensity in the VIS, SMN, CON, DMN, FPN, AUD, DAN and HIPP during encoding all showed significant correlations with the memory scores, whereas during retrieval, only that of the CON, DMN and FPN displayed significant correlations with the memory scores (r > 0.35, p < 0.01; Figure 6a).

**Figure 6.**
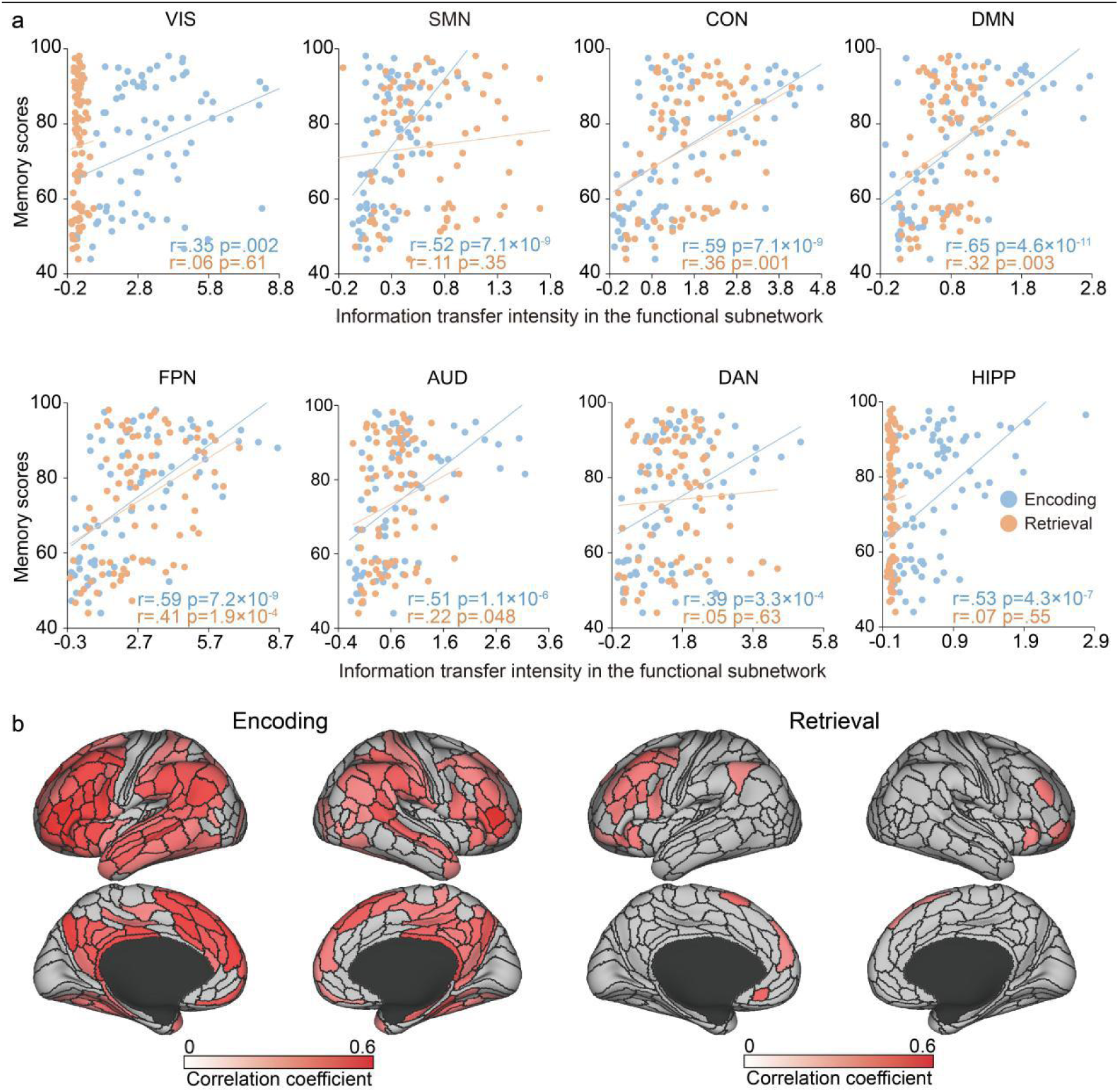
The correlation between information transfer intensity and memory performance during encoding and retrieval. Memory performance was computed by the confidence rating in the episodic memory task. **a** The correlation at the subnetwork level. The horizontal axis represents the average information transfer intensity of each functional subnetwork. **b** The correlation at the regionwise level. All reported subnetworks (regions) were statistically significant at p < 0.01.

Furthermore, more widely distributed regions showed significant correlations with the memory scores during encoding (193 regions) than during retrieval (27 regions) at the regional level (see Figure 6b; for the correlation between each transfer and memory performance, see Figure 6—Supplementary figure 1). Most of the regions within the FPN revealed significant correlations during both encoding and retrieval (Pearson’s r>0.29, p<0.01). In addition, most regions in the CON, DMN and HIPP (parahippocampal and hippocampal) and fewer regions in the VIS (mainly in the ventral stream of the visual cortex), SMN (somatosensory and motor cortex, premotor cortex, paracentral lobule and midcingulate cortex), AUD (mainly in the auditory association cortex) and DAN (mainly in the superior parietal cortex) showed predictability during encoding (Pearson’s r>0.29, p<0.01). Compared with encoding, only fewer regions in the CON and DMN had this characteristic (Pearson’s r>0.29, p<0.01) during retrieval (Figure 6b). This result indicates that information transfer had a significant effect on memory performance, which is more significant in the encoding process.

### Building a prediction model based on the information transfer network

The above results showed that information transfer is closely related to memory performance. Hence, we used the information transfer network as the main feature and combined connectome-based predictive modeling to measure the predictive power of memory performance (see materials and methods for details). Figure 7 shows the correlation between observed memory scores and predicted memory scores. Three different feature selection methods (Pearson correlation, Spearman correlation and robust regression) all showed predictive power for memory performance using the information transfer network during both encoding (Pearson’s r=0.60, p=4.4×10^-9^; r=0.61, p=1.8×10^-9^; r=0.60, p=4.6×10^-9,^ respectively) and retrieval (Pearson’s r=0.31; p=0.005; r=0.38, p=5.15×10^-4^; r=0.34, p=0.002, respectively). Moreover, the information transfer network had higher predictive power during encoding than during retrieval (Steiger’s z = 2.53, p-value = 0.01; z = 2.18, p = 0.03; z = 2.35, p = 0.02, respectively).

**Figure 7.**
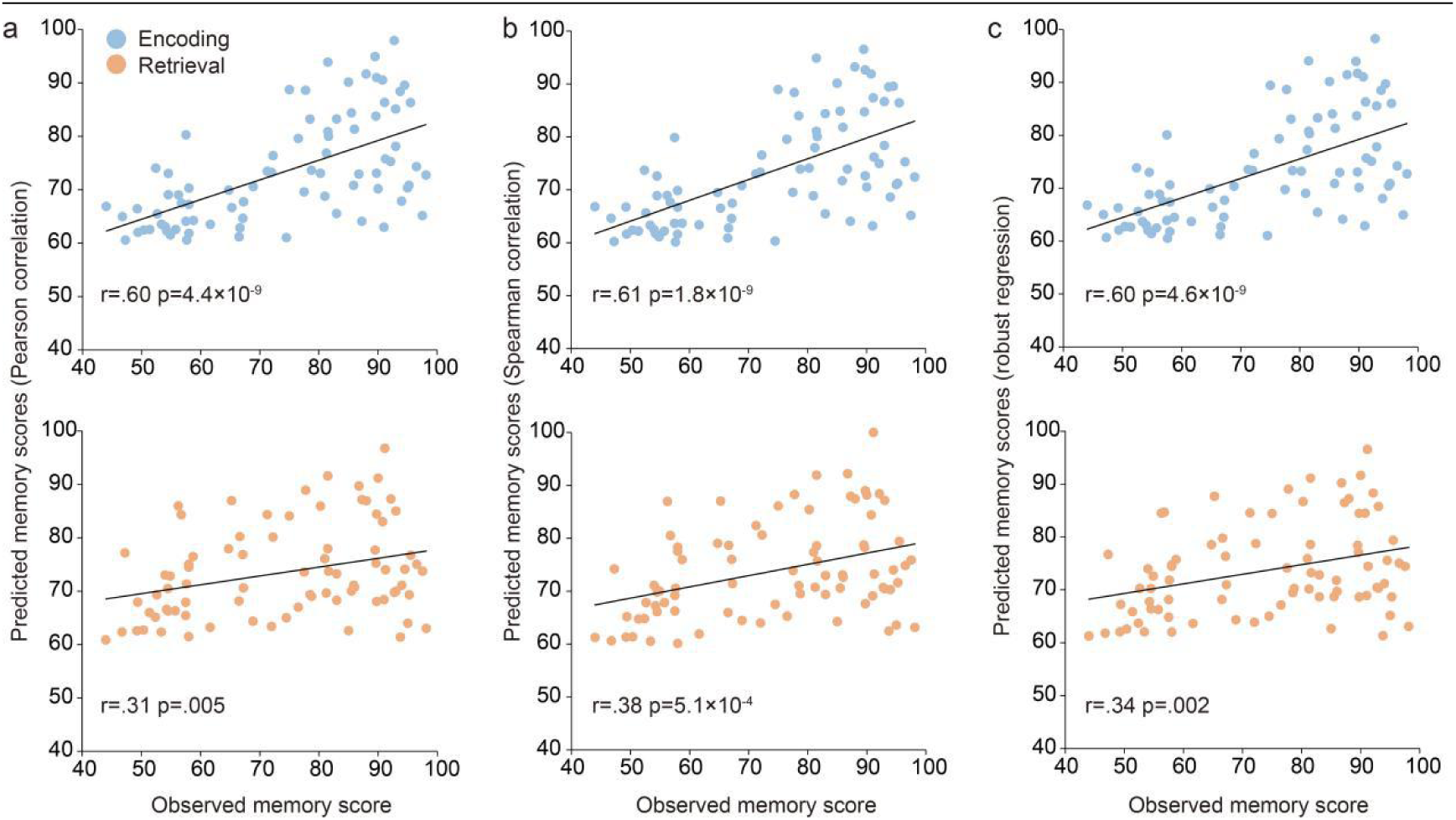
Correlation between predicted and observed memory scores. A region-to-region information transfer network was employed to predict episodic memory performance. **a**, **b**, and **c** represent three different feature selection methods: Pearson correlation, Spearman correlation, and robust regression (edge-selection threshold of p < 0.01).

To confirm the advantage of information transfer, we also used the same model, with tasking-state FC and resting-state FC as the features, to predict memory scores. The results showed that the predictive ability of information transfer was significantly stronger than that of task-state FC during both encoding (Steiger’s z = -2.30, p-value = 0.02; Steiger’s z = -2.50, p-value = 0.013; Steiger’s -z = 2.22, p-value = 0.02, respectively) and retrieval (the task-state FC network during retrieval could not accurately predict memory, p>0.01) (Figure 7—Supplementary figure 1). The predictive ability of resting-state FC is similar to that of information transfer during retrieval (Pearson: Steiger’s z = -0.026, p-value = 0.97; Spearman: Steiger’s z = 0.31, p-value = 0.76) but is obviously weaker than that during encoding (Pearson: Steiger’s z = -2.59, p-value = 0.0095; Spearman: Steiger’s z = -1.47, p-value = 0.048) (Figure 7—Supplementary figure 2). Therefore, these results indicate that information transfer could act as an effective indicator of episodic memory performance.

### The mediating effect of information transfer

Given that information transfer was significantly correlated with both task-evoked activation and memory performance, we further explored the relationship among information transfer, task-evoked activation, and memory performance using mediation analysis. We found that memory performance was more correlated with information transfer intensity than with task-evoked activation (Figure 5 and Figure5—Supplementary figure 2). Information transfer may thus have a direct effect on memory performance. We next selected the regions shown in Figure 5 for mediation analysis (the positively and negatively correlated regions were analyzed separately) and regarded the information transfers of these regions as the mediators. Figure 8 shows the relationships between information transfer intensity, task-evoked activation and memory performance. We observed that during encoding, the averaged activation of positively correlated regions had significant direct effects on memory performance (path c’: β=14.52, SE=7.26, t=2.00, p=0.049) and had an indirect effect of information transfer on memory performance (path ab: β=19.10, SE=5.22, t=3.66, p<0.001). In contrast, the averaged activation of negatively correlated regions only had an indirect effect on memory performance (path ab:β=-18.34, SE=5.47, t=-3.35, p<0.001). During retrieval, the averaged activation of both positive and negative regions had no direct effect on memory performance (respective path c’: β = 9.09, SE = 11.49, t = 0.79, p = 0.43; β=22.01, SE=11.28, t=1.95, p=0.06) but had an indirect effect on memory performance through information transfer (respective path ab: β=17.38, SE=6.48, t=2.69, p=0.007; β=-10.17, SE=4.98, t=-2.04, p=0.04). These results indicate that information transfer has an incomplete mediating effect on brain activation and memory during encoding but a complete mediating effect during retrieval.

**Figure 8.**
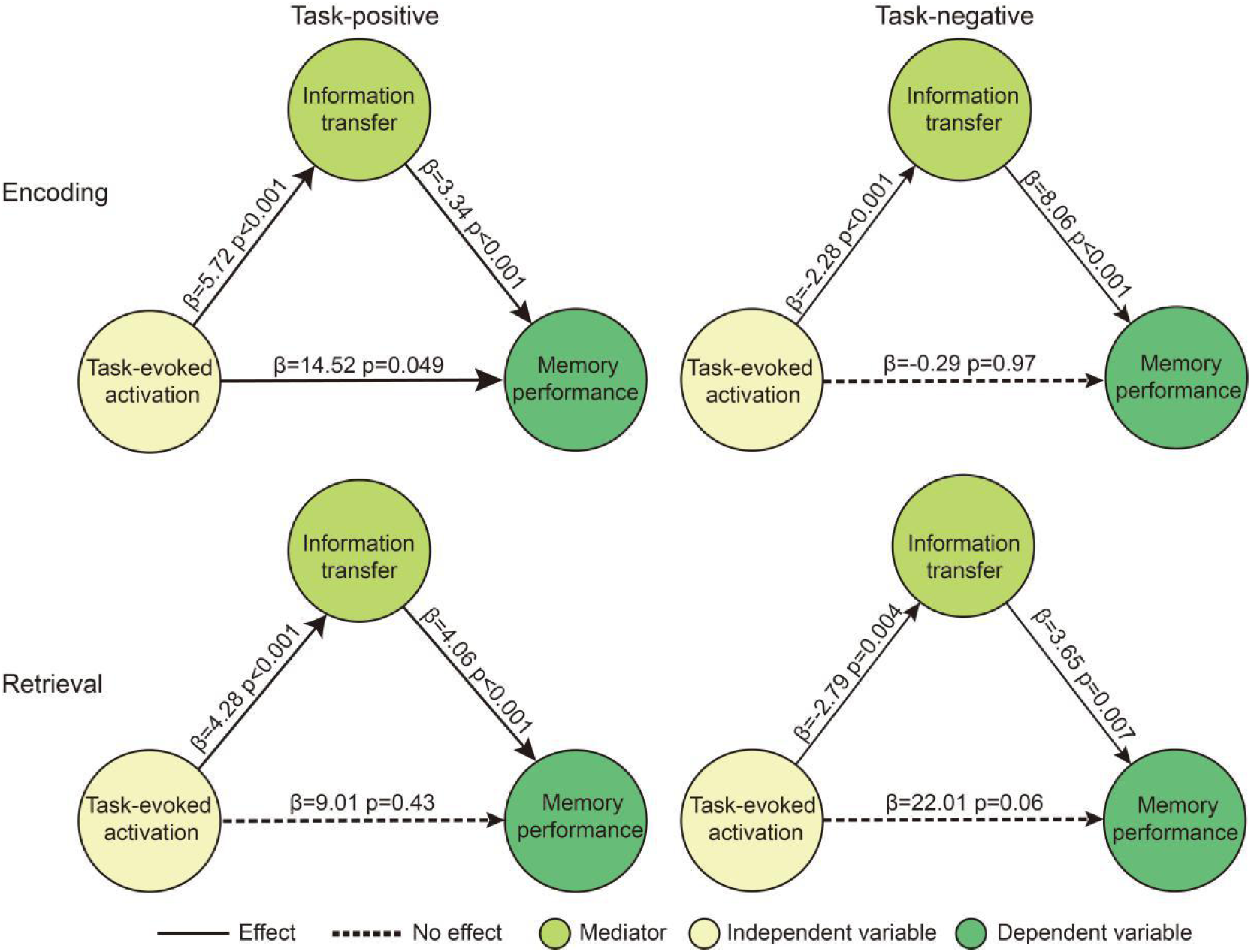
Mediation analysis between task-evoked activation, information transfer intensity and memory performance. The significance of the mediating effect was measured by a threshold of p < 0.05 of the Sobel test. An incomplete mediation effect was found in TP regions (indirect effect: β=19.10, SE=5.22, t=3.66, P<0.001; direct effect:β=14.52, SE=7.26, t=2.00, P=0.049), and a complete mediation effect was found in TN regions (indirect effect: β=-18.34, SE=5.47, t=-3.35, P<0.001; direct effect: β = 9.09, SE = 11.49, t = 0.79, p = 0.43) during encoding. Complete mediation effects were evident in TP regions (indirect effect: β=17.38, SE=6.48, t=2.69, P=0.007; direct effect: β = 9.09, SE = 11.49, t = 0.79, p = 0.43) and TN regions (indirect effect: β=-10.17, SE=4.98, t=-2.04, P=0.04; direct effect: β=22.01, SE=11.28, t=1.95, P=0.06) during retrieval. (TP: task positive, TN: task negative).

### SCs and information transfer

Although brain function based on structural and regional communication also requires some material support, the structural foundation underlying functional information transfer is unclear.

We found that the intensity of direct transfers was significantly higher than that of indirect transfers (p<0.001, details see Supplementary information), and direct transfer intensity increased as the threshold of defining an SC became stricter (Figure 9). However, the correlations between these two structural types of transfer and memory performance were not significantly different (Table 1). These results indicate that an SC has an effect on information transfer intensity, which may play a role in transmitting information, whereas both indirect and direct transfer have important roles in episodic memory.

**Figure 9.**
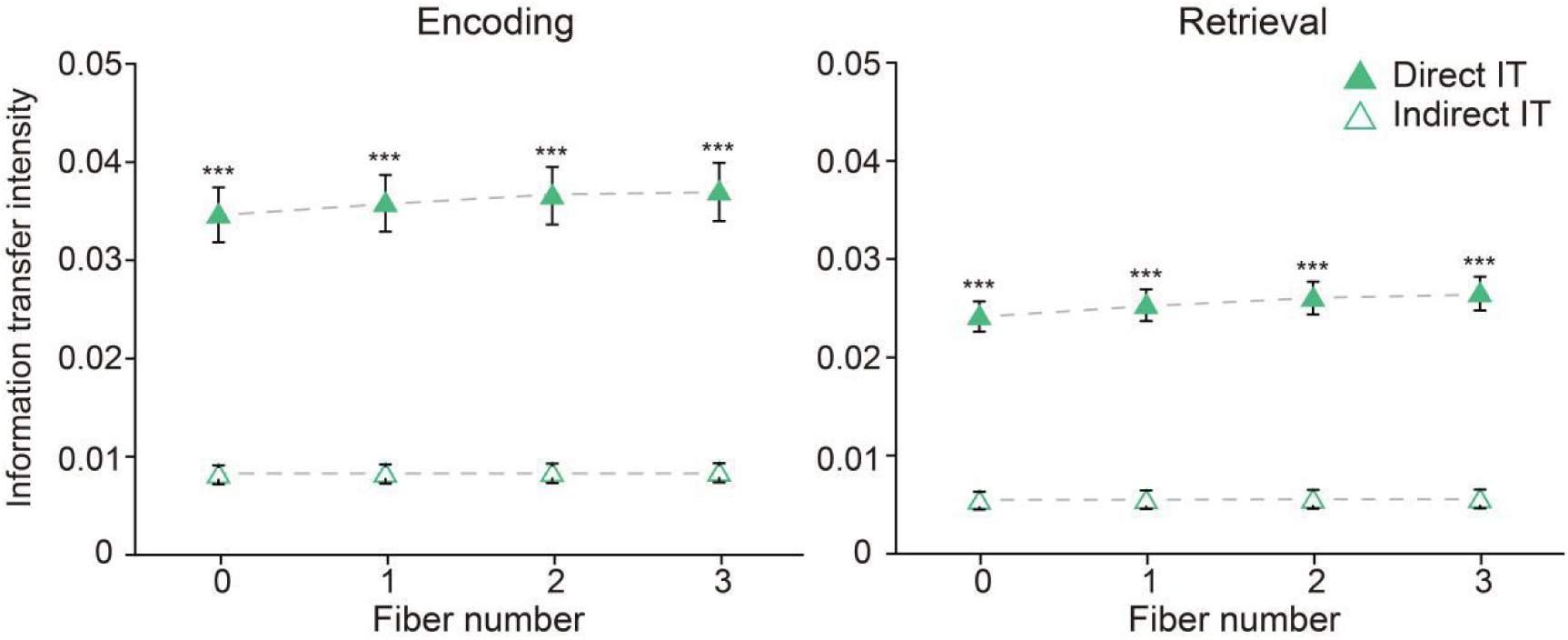
Influence of SCs on information transfer intensity. Information transfer is divided into direct or indirect transfers according to whether there were SCs between regions (the setting threshold of the fiber number was from 0 to 3). *** represent p < 0.001, IT = information transfer.

**Table 1.** Effect of direct vs indirect information transfer on memory

| **Table 1** Effect of direct vs indirect information transfer on memory |  |  |  |  |
| --- | --- | --- | --- | --- |
| **Paradigm** | **FN** | **Direct IT** | **Indirect IT** | **Steiger** |
| Encoding | 0 | r=0.601 p=7.14e-9 | r=0.552 p=1.85e-7 | z=1.455 p=0.145 |
|  | 1 | r=0.606 p=5.09e-9 | r=0.553 p=1.79e-7 | z=1.603 p=0.108 |
|  | 2 | r=0.614 p=2.88e-9 | r=0.553 p=1.74e-7 | z=1.88 p=0.059 |
|  | 3 | r=0.613 p=3.01e-9 | r=0.554 p=1.64e-7 | z=1.49 p=0.136 |
| Retrieval | 0 | r=0.271 p=0.016 | r=0.256 p=0.024 | z=0.341 p=0.733 |
|  | 1 | r=0.262 p=0.021 | r=0.257 p=0.024 | z=0.115 p=0.908 |
|  | 2 | r=0.269 p=0.017 | r=0.257 p=0.024 | z=0.271 p=0.786 |
|  | 3 | r=0.267 p=0.018 | r=0.257 p=0.024 | z=0.299 p=0.765 |
| Direct and indirect transfer were divided according to the set threshold of FN. r values refer to the correlation coefficient between transfer and memory performance, and z values refer to the difference in correlation coefficient. SC=structural connection, FN= fiber number, IT=information transfer. |  |  |  |  |

### Validation

We validated our results regarding information transfer in three aspects: its role in the pathological state of an individual, the correlation of information transfer with brain state changes, and information transfer in artificial neural networks (ANNs).

First, given that prior studies suggest that patients with bipolar disorder (BD) have abnormal cognitive function, including episodic memory (Bora, Yucel et al. 2009, Hibar, Westlye et al. 2018), we therefore expected that information transfer could reflect episodic memory performance and serve as an indicator of episodic memory function. We compared the information transfer and task-evoked activation between healthy people and patients with BD. The results showed that there was no difference in task-evoked activation of each network between the two groups (Figure 10a), but there was a significant difference in information transfer intensity (p<0.05 with Bonferroni correction, Figure 10b). This result suggests that information transfer was more sensitive than the traditional task-evoked activation approach for detecting BD dysfunction.

**Figure 10.**
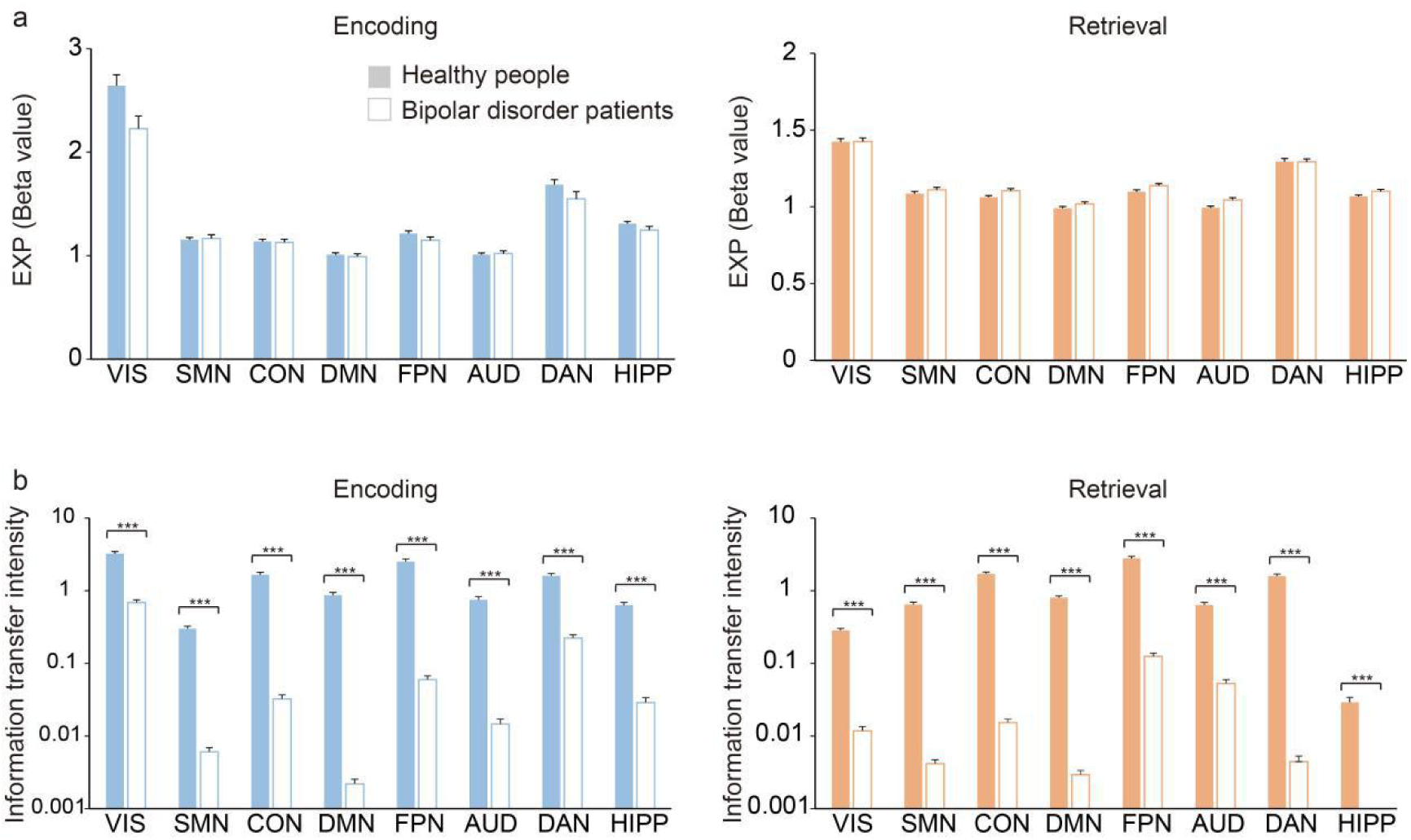
The validation of the results through a comparison between BD patients and healthy people. **a** Comparison of beta values. None of the functional subnetworks showed significant differences (p<0.05 with Bonferroni correction). **b** Comparison of information transfer intensity. Each functional subnetwork showed significant differences (p<0.05 with Bonferroni correction). EXP represents the exponential function with base e. *** represents p<0.001 with Bonferroni correction; error bars indicate SE of estimates.

Second, we also expected that our results could accurately reflect changes in one’s brain state. We used blood oxygen level-dependent (BOLD) signals at the beginning and end of each trial to calculate the changes in brain state (see materials and methods for details). We found that information transfer intensity was significantly correlated with changes in the brain state of most subjects. In the ‘memory’, information transfer intensity could characterize the changes in the brain state of 87.5% of healthy people and 91% BD patients. Regarding the ‘correct’ information transfer outcome, 74% of healthy people and 71% of BD patients showed this effect, and in the ‘incorrect’ information transfer outcome, 68% of healthy people and 76% of BD patients showed this phenomenon (Figure 10—Supplementary figure 1). This finding suggested that our results could describe changes in the brain state during epsiodic memory.

Third, because ANNs are created based on brain networks, we built an ANN model to simply simulate the working process of the brain, including signal encoding, information transfer, and signal decoding (see materials and methods for details). The results show that output accuracy was higher when the effective transfers (Figure 2a) were retained (encoding: 88.03%; retrieval: 95.06%) than when the ineffective transfers were retained (encoding: 73.21% ± 2.71%; retrieval: 82.93% ± 3.36%) (Figure 10—Supplementary figure 2). This finding showed that our results were also repeatable in an ANN.

## DISCUSSION

This work examined information transfer patterns during encoding and retrieval, aiming to provide a more intuitive understanding of the brain communication mechanism underlying episodic memory. We found that the CON, DMN, FPN and DAN were engaged in information transfers of episodic memory, and the transfer between the CON and FPN was most significant in both encoding and retrieval. The VIS and HIPP were more engaged in information transfer during encoding, whereas the SMN was more engaged in information transfer during retrieval. Information transfer intensity in the CON, DMN and FPN was related to task-evoked activation. Information transfer was shown to accurately encode information related to memory ability and demonstrated considerable predictive power for memory performance. Furthermore, information transfer could mediate the relationship between task-evoked activation and memory performance. In addition, SCs were shown to be a carrier of information during transfer. Finally, from multiple perspectives, our study verifies that a combination of resting-state FC and task-evoked activation reflects regional communication of episodic memory more accurately.

### Commonality in information transfer between encoding and retrieval

We found that both encoding and retrieval processes recruited transfers within the HIPP, CON, FPN, DAN, VIS and DMN. The CON, DMN, FPN and DAN were the most recruited brain networks during episodic memory. This result was also observed in a multitask paradigm (Ito, Kulkarni et al. 2017). A large number of studies have indicated the importance of these networks in episodic memory. Notably, the HIPP has been recognized as an essential part of memory formation (Moscovitch, Cabeza et al. 2016, Sekeres, Winocur et al. 2018). Also, the FPN has been shown to support cognitive control and integrate memory with attentional resources allocation (R Nathan, Jorge et al. 2013, Dixon, De La Vega et al. 2017). The CON has been revealed to be primarily involved in task maintenance (Cabeza, Ciaramelli et al. 2008, Carlo, Maurizio et al. 2014, Rugg and King 2017) and the DMN has been associated with internally directed cognition (Chai, Ofen et al. 2014, Murphy, Jefferies et al. 2018). Pertinently, the DAN has been associated with the conversion, search and detection of spatial attention (Ciaramelli, Grady et al. 2008, Sestieri, Shulman et al. 2017). Moreover, the VIS has been shown to receive visual information inputs and send them to a higher-level cognitive network [(Bar, Kassam et al. 2006, Wen, Shi et al. 2018). These findings demonstrate that the neural resources of cognitive control, attentive control, memory storage and visual reception systems are all recruited in information transfer processes of episodic memory.

It has been confirmed that the FPN and CON functionally interact (Power and Petersen 2013) and belong to multiple demanding cognitive systems. Information transfer from the FPN to CON was shown to be significant during multitask execution (Ito, Kulkarni et al. 2017). The FPN has been identified as being responsible for top-down modulation during preparatory attention and orienting within memory processes (Gui, Qi et al. 2012, Wallis, Stokes et al. 2015). In contrast, the CON was found to play a more downstream role in cognitive control and provides stable set maintenance (Sestieri, Corbetta et al. 2014, Wallis, Stokes et al. 2015). The FPN sends instructions to the CON, and then the CON gives feedback to the FPN; thus, the high level of information transfer between them imply an important mode during episodic memory. Moreover, previous studies have substantiated that the DMN and DAN have a reciprocal relationship—the activation of one network inhibiting the another—and the two networks are modulated by the CON and FPN as they flexibly couple with either network depending on the task domain (Carlo, Maurizio et al. 2014, Kucyi, Daitch et al. 2020). Therefore, this evidence suggests that encoding and retrieval also depend on information transfer between the control system and the attention system for switching and maintaining attention. Notably, these conclusions are supported by other cognitive task studies about information communication (Krienen, Yeo et al. 2014, Avenakoenigsberger, Misic et al. 2018, Toker and Sommer 2019).

### Differences in information transfer patterns between encoding and retrieval

The comparisons of information transfer patterns between encoding and retrieval showed some differences between these two processes. Specifically, the HIPP was engaged in more transfers during encoding than retrieval and transferred information via the VIS, DMN and FPN only during encoding. This may be due to the specific role of the hippocampus and parahippocampus in ‘recollection’ but not in ‘familiarity’ (Tanaka, Pevzner et al. 2014). The subjects only needed to discriminate between studied and nonstudied items rather than retrieve specific information about each item in the retrieval task, whereas the HIPP processes a considerable amount of external information in encoding, especially in relational processing (Moscovitch, Cabeza et al. 2016, Sekeres, Winocur et al. 2018). Thus, it is possible that HIPP was less involved in retrieval and thereby did not induce significant information transfer compared with encoding tasks. Furthermore, HIPP also transferred information with the VIS, DMN and FPN only during encoding. Studies have suggested that HIPP plays an important role in visual representation and perception (Martin, Douglas et al. 2018). Thus, the transfers between the VIS and HIPP were mainly used to represent the objects of interest (for example, the meaning of words and images), thereby enhancing memory, whereas in retrieval tasks, the subjects only needed to determine the object’s familiarity, which was less dependent on visual discrimination. Moreover, the DMN provided a bridge linking cortical areas with the HIPP, though which information reached the HIPP via the DMN (Rolls 2019). Thus, this result indicated that information transfer between the HIPP with DMN and FPN was an important basis for memory formation.

The VIS was also involved in more information transfer during encoding. When a ‘novel’ target appeared, the VIS extracted its features through massive transfers and transmitting the information to the ‘advanced’ network, which in turn transmitted instructions to the VIS (Bar, Kassam et al. 2006). During retrieval, some encoded representations in the VIS were reinstated in the FPN (Xiao, Dong et al. 2017). It has been suggested that when “familiar” objects appear, the brain only needs to extract some valuable features instead of representing them completely (Armann, Jenkins et al. 2016). Thus, our result suggests that the information transfer of the VIS is determined by the number of features.

In contrast to the HIPP and VIS, we found that the superior parietal cortex of the DAN was involved in greater information transfer during retrieval. This cortical section has been shown to be maximally engaged in top-down attentional control of prior information when ‘familiar’ objects appear (Ciaramelli, Grady et al. 2008, Sestieri, Shulman et al. 2017). Therefore, the superior parietal cortex induced a higher level of information transfer to focus attention on the ‘familiar’ object during retrieval.

### Correlation between information transfer and brain activation

Our analyses showed that the regions with positive correlations between information transfer and activation patterns were mostly located in the FPN. Notably, activation of brain regions has been shown to reflect the metabolic level of neurons (Magistretti and Allaman 2018). This finding might imply that the FPN could flexibly control the information transfer pattern to alter the interaction with other networks during episodic memory. In support of this proposal, the FPN was shown to be a flexible hub in cognitive control that adapts to various task demands by flexibly interacting with a variety of functionally specialized networks (Cole, Pathak et al. 2011, Cole, Reynolds et al. 2013, Avenakoenigsberger, Misic et al. 2018). Thus, these findings propose that the FPN could serve as an initiator of the information transfer process in episodic memory. During encoding, many regions in the CON also showed positive correlations. The CON was found to be functionally strongly related to the FPN. Thus, the CON also was shown to have flexible characteristics in information transfer. During retrieval, the information transfer pattern of the CON was stable, which indicated that the ‘familiarity’ mechanism is relatively simple and did not need much adjustment. Additionally, the regions with negative correlations were mainly located in the posterior cingulate and the inferior parietal cortex, which were involved in internally oriented processing (Murphy, Jefferies et al. 2018) and bottom-up attention to retrieval outputs (Sestieri, Shulman et al. 2017). Studies have revealed that these regions belong to a task-negative network that deactivates episodic memory (Kim, Daselaar et al. 2010, Medaglia, Satterthwaite et al. 2018). Therefore, this result confirmed that although deactivated during the memory task, the DMN network has the ability to communicate with other regions and can flexibly adjust information transfer during both encoding and retrieval.

### Relationship between information transfer and memory performance

Similar to the correlation with activation, we found that many information transfers of the FPN, CON and DMN strongly affected memory performance in both encoding and retrieval. As mentioned above, the CON and FPN are hubs of brain function. Combining their information transfer patterns, the results indicated that the information transfer of control-related networks had powerful functions to integrate and reconstruct episodic details. Thus, the CON and FPN were found to be the most important components in the episodic memory mechanism, and information transfer was shown to be an essential operation mode. The DMN has also been shown to be critical for cognition and mainly supports internal attentive control (Cabeza, Ciaramelli et al. 2008, Chai, Ofen et al. 2014, Murphy, Jefferies et al. 2018). Our results suggested that the information transfer of the DMN may have inhibited internal attention, paying attention to external stimuli during encoding, while arousing internal attention during retrieval. Thus, information transfer of the DMN contributed to internal attention allocation and control both by promotion and inhibition mechanisms.

In addition, information transfer of the HIPP significantly affected memory performance during encoding. Studies have suggested the importance of the HIPP from multiple the perspectives of lesions, regional activity, FC and graph theory (Sekeres, Winocur et al. 2018). The present study confirmed from a new perspective that information transfer of the HIPP is also an indispensable component during encoding. Furthermore, we found that information transfer of the HIPP had no effect on memory performance during retrieval. It may be that HIPP is involved in weaker information transfer in ‘familiarity’.

Interestingly, our result also suggested that the information transfer of the SMN had an effect on memory performance during encoding. The premotor cortex has been shown to be involved in the maintenance of visuospatial attention and response delay (Owen 2005), and the midcingulate cortex could be related to an additional attention process mediating information between the VIS and FPN (Roy, Jamison et al. 2017). This finding suggests that although there were fewer transfers in the SMN during encoding, the functional role of the SMN remains crucial for cognitive control and one’s response to a stimulus. Successful retrieval mainly depends on internal attention but depends less on the response to external stimuli (Kim, Daselaar et al. 2010). Therefore, during retrieval, information transfer in the SMN may have no such characteristic. Additionally, the VIS, DAN and AUD also showed significant correlations only during encoding. The VIS involved a considerable amount of information transfers to characterize external objects. The transfers in the DAN were used to maintain external attention. Also, the retrieval task was mainly guided by internal processing. Therefore, these areas played more obvious roles in encoding. In AUD, the primary auditory cortex mainly processes auditory information, whereas the auditory connection cortex and temporoparietal junction mainly encode semantics and construct words. No auditory stimuli were used in the present study, and information transfer in the AUD was used to understand the meaning of words during encoding.

Using connectome-based predictive modeling (Shen, Finn et al. 2017), we further demonstrated that information transfer was a more powerful predictor for memory performance than tasking- and resting-state FC. Thus, the region-to-region information transfer network could serve as a holistic neural index for predicting episodic memory ability.

### A mediating role of information transfer

Of note, information transfer of the FPN, CON and DMN showed significant correlations with both task-evoked activation and memory performance. Memory performance was affected more by information transfers than task-evoked activation. Task-evoked activation has been shown to reflect the metabolism of neurons, which provides energy for signal generation (Hahn, Poncealvarez et al. 2019). Therefore, this finding confirmed that information transfer could be the product of task activation, thereby it would have a more direct effect on memory function. Then, by using the mediate analysis, our result suggested that the information transfers of the FPN, CON and DMN (figure 4a) could directly affect memory storage and integration during encoding. In contrast, task-evoked activation in the FPN and CON (figure 4b) indirectly affected memory retrieval through the intermediate effect of information transfers.

### Influence of structural connectivities in information transfer

We also explored the influence of SCs on information transfer. Previous studies have suggested that SCs affect brain function (Memel, Wank et al. 2020) but have not found an obvious link between SC and FC with respect to episodic memory (Suarez, Markello et al. 2020). However, we found that SCs induced stronger information transfer, and this characteristic increased with the number of fiber bundles. This result suggested that fiber bundles are carriers of functional information.

It is important that we discuss the generalizability and reliability of the results from three aspects: psychiatric illness, brain activity status and simulation. We confirmed that information transfer may be used as a detection index of brain diseases with episodic memory impairment and could be used as a predictor of brain state changes; moreover, information transfer even has certain advantages in establishing artificial brain models. Thus, the findings in the current research could be used to describe the information transfer pattern in episodic memory.

Collectively, this was the first study to apply information transfer mapping to investigate the brain network mechanism of episodic memory. The results provided new insight into the brain communication mechanism of episodic memory and confirmed the advantages of a combined analysis of task-related and resting-state fMRI data by a novel approach comprising activity flow mapping in explaining memory function.

## Materials and methods

### Participants

The neuroimaging dataset was shared from the UCLA Consortium for Neuropsychiatric Phenomics LA5c Study and approved by the UCLA Institutional Review Board (Poldrack, Congdon et al. 2016). 139 subjects with no relevant medical or psychiatric information participated in the experiment. 80 subjects (females=37, mean age=30.03 years, SD=8.29, range=21 - 49), collected the both resting-state and tasking-state fMRI data with high quality, were include in our analyses. The data were obtained via the public database OpenfMRI.

### Imaging acquisition

Whole-brain imaging data were acquired on one of two 3 T Siemens Trio scanners (Siemens version Syngo MR B15 and B17). Resting-state and tasking-state fMRI data were collected using a T2-weighted echoplanar imaging (EPI) sequence with the following parameters: repetition time (TR) = 2 s, echo time (TE) = 30 ms, flip angle = 90°, slice thickness = 4 mm, slices = 34, matrix = 64 × 64, and field of view (FOV) = 192 mm. In addition, a high-resolution anatomical magnetization-prepared rapid acquisition with gradient echo (MPRAGE) scan was collected. The parameters for MPRAGE were as follows: TR = 1.9 s, TE = 2.26 ms, matrix = 256 × 256, FOV = 250 mm, sagittal plane, slice thickness = 1 mm, and slices = 176.

### Behavioral paradigm

We used a paired associate memory task, which is a classic paradigm of episodic memory (Figure 1a). In the task, two scans were performed to assess declarative memory encoding and retrieval. Memory encoding included 40 “memory” trials and 24 control trials. During “memory” trials, two screens each displayed a word for 1 s, then line drawings of two objects that matched each word appeared above the word; the word and the object were presented together for an additional 3 s. One of the objects was drawn in black and white, and one object was drawn using a single color. For control trials, pairs of scrambled stimuli, in which one screen was black and white and another was colored, lasted for 2 s and participants selected which side the colored object was on by button press. Subjects were instructed to remember the objects and their relationship with the word. The retrieval task included 40 ‘correct’ trials, 40 ‘incorrect’ trials and 24 control trials. The retrieval task required the subjects to look at a pair of objects and rate their confidence in their memory of the pairing. There were 4 possible response options ranging from “sure correct” to “sure incorrect”, allowing the responses to be analyzed as a spectrum or binarized into yes/no type responses. During control trials, one side of the screen displayed one of the four retrieval confidence response options (“sure correct”, “maybe correct”, “maybe incorrect”, or “sure incorrect”). On the other side of the screen, “xxxx” was displayed. Subjects were asked to press the button (1-4) that corresponded to the response option displayed (for example, a response of 1 if “sure correct” appeared. In the 40 correct trials, items were shown paired as they had been at encoding. During the 40 incorrect trials, items were shown paired differently than they were at encoding; some objects were the same but were just paired incorrectly. At the end of the task, memory performance was evaluated according to the accuracy of retrieval.

### Memory score calculation

The subjects were graded according to the accuracy of their rating confidence regarding the retrieval process. There were 4 types of response answers ranging from “completely right choice” (choosing “sure correct” in “correct” trials or “sure incorrect” in “incorrect” trials) to “completely wrong choice” (choosing “sure correct” in “incorrect” trials or “sure incorrect” in “correct” trials) corresponding to 4 types of scores: 5, 3, -3 and -5. Then, the scores of each trial were summed, and the total score (0 to 100 scores) was normalized, which was used to represent memory performance.

### fMRI preprocessing

Resting-state and task-state fMRI data were preprocessed using SPM12 by performing the following steps: (1) slice time correction, (2) head motion correction with alignment to the minimum outlier in the time series, (3) removal of the effect of nuisance covariates, (4) registration with T1-weighted image unified segmentation and spatial normalization to the MNI template, (5) Gaussian filter smoothing, and (6) low-frequency filtering processing.

In addition, using a standard fMRI general linear model (GLM) combining task onsets with the SPM canonical hemodynamic response function (HRF) and adding 12 motor parameters (6 for rigid-body translation and 6 for rotation plus their derivatives), ventricle and white matter time series in GLM were used to analyze preprocessed task images. The GLM produced trial-by-trial task-related beta activity at the voxelwise level, which was used to indicate task-evoked activation.

### Brain region parcellation

Voxels were divided into regions according to the brain atlas established by Glasser et al. (Glasser, Coalson et al. 2016). The Glasser atlas divided the brain into 22 cortex sections and then subdivided them into 360 regions. The cortex section names all came from the Glasser atlas in the present study. Then, the regions were assigned to the functional subnetworks using the generalized Louvain method. The present study focused on eight major networks: the VIS, SMN, CON, DMN, FPN, AUD, DAN, and HIPP (Figure 1b).

### Functional connectivity estimation

Multiple linear regression was used to estimate resting-state FC at the voxelwise level. To estimate FC for a given node, standard linear regression was used to fit the time series of all other nodes as predictors (i.e., regressors) of the target nodes. Using ordinary least squares regression, whole-brain FC estimate values were calculated by obtaining the regression coefficients from the equation.

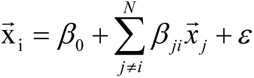

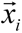 is the time series in voxel 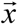, *β*_0_ is the y-intercept of the regression model, *β_ij_* is the FC coefficient for the *j*th regression/voxel (the element in the *j*th row and the *i*th column in the FC adjacency matrix), and *ε* is the residual error of the regression model.

### Region-to-region information transfer mapping

The activity flow mapping procedure unified both biophysical and computational mechanisms into a single information-theoretic framework to allow for an investigation into the transfer of task-related information between pairs of brain regions using voxel activation patterns (Cole, Ito et al. 2016). This proved that resting-state FCs describe the large-scale architecture of information communication in the human brain and demonstrate the relevance of resting-state network connectivities to cognitive information processing.

The activity of the voxel of a held-out target region was predicted based on the voxel within a source region (Figure 1c). Region-to-region activity flow mapping between region A and region B was defined as:

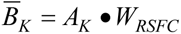

where 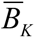 corresponds to the predicted activation pattern vector for target region B, *A_K_* corresponds to the activation pattern vector of region A (i.e., the source region), *W_RSFC_* corresponds to the voxel-to-voxel resting-state FC between regions A and B, and the operator • refers to the dot product. This formulation mapped activation patterns in the spatial dimension of one region to the spatial dimension of another.

Then, information transfer mapping was used to quantify how much information transfer occurred between two regions (Figure 1c). Briefly, information transfer mapping comprised three steps (Ito, Kulkarni et al. 2017): (1) region-to-region activity flow mapping; (2) cross-validated representational similarity analysis between predicted activation patterns and actual, held-out activation patterns; and (3) information classification/decoding by computing the difference between matched condition similarities and mismatched condition similarities. ITE_AB_ denotes the information transfer from region A to region B and is defined as:

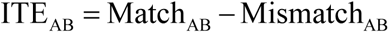

where Match*_AB_* and Mismatch*_AB_* correspond to the averaged Pearson rank correlation for matched and mismatched conditions using source region A, respectively. Match_AB_ and Mismatch_AB_ were defined as:

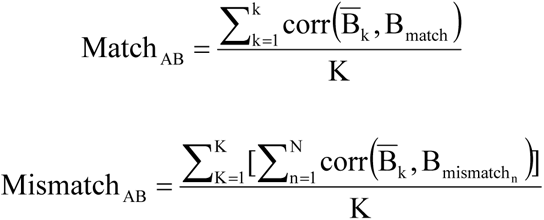

where k corresponds to the total number of trials; corr corresponds to a Fisher–Z transformed Pearson’s rank correlation between two vectors; B_K_ is defined as the predicted activation pattern in target region B (using the activation pattern of region A) for trial K; B_match_ is defined as the condition prototype (obtained by averaging across trials of the same condition, holding out a trials) of the actual activation pattern for K in target region B, which the condition of the trial matches; and B_mismatch_ is defined as the condition prototypes for which the condition of the trials does not match.

This formulation quantified how much “information transfer” occurred between two regions by comparing the predicted activation pattern in the target region to the actual activation pattern in the target region across all cross-validation folds.

Statistical tests were performed using a group one-tailed t-test for every pairwise transfer. The transfers significantly greater than 0 (p<0.05 with FDR correction) in the statistical tests were deemed effective; otherwise, they were considered ineffective. Any significant deviation above 0 indicated a significantly higher matched correlation compared to the mismatched correlation between predicted-to-actual activation patterns. Any information estimate that was not significantly greater than 0 indicated that the predicted-to-actual similarity was at chance level.

Then, the percentage of significant network-to-network transfers was calculated (the number of significant transfers divided by the number of possible network-to-network transfers), which could provide better visualization and allow for the evaluation of region-to-region transmission mapping that may have been affected by the underlying network organization (if the transfers between all pairs of regions in network affiliations were less than 2, it was determined that there was no effective transfer).

### Regional information transfer analysis

The regional information transfer percentages were obtained by taking the number of successful transfers from and to a region and dividing it by the total number of possible transfers (i.e., 359×2 other regions). The significant transfer percentage was calculated for each region to intuitively represent the number of transfers sent and received by each region and to visualize the anatomical locations of the regions.

Regional information transfer intensity was defined as the weighted sum of significant information transfers sent and received by a region, indicating its efficiency of processing information and sensitivity to information. A paired t-test was used to compare the information transfer intensity between encoding and retrieval for each subnetwork (region) (significance at p < 0.05 with).

The Pearson correlation coefficient was used to measure the relationship between regional information transfer intensity and the regional beta value and to estimate the effect of the information transfer of each network (region) on memory scores (significance at p<0.01).

### Prediction model establishment

Connectome-based predictive modeling was employed to estimate the episodic memory ability of subjects (Shen, Finn et al. 2017). First, correlation analysis (Pearson correlation, Spearman correlation or robust regression) between each edge in the information transfer matrices and memory scores was performed across subjects. Second, a threshold was applied to the matrix that only retained edges that were significantly and positively correlated with behavior (p < 0.01); the edges from the information transfer matrix of one subject were summed, which represented the strength of the whole-brain information transfer network. Next, a leave-one-out cross-validation procedure was employed. In each set of n − 1 subjects, predictive networks were defined as described above. Simple linear models were then constructed to relate information transfer network strength to memory scores during encoding and retrieval in these subjects. Finally, these models were used to predict the left-out subject’s memory score. Pearson correlations between observed and predicted memory scores were used to assess predictive power (significance at p<0.01). Steiger Z tests were used to compare two overlapping correlations (significance at p<0.05).

### Mediation analysis

Mediation analysis was used to examine whether there was a mediating effect between task activation, information transfer intensity, and memory performance (Zhao, Jr et al. 2010). First, regions where the intensity of information transfer was significantly correlated with task-evoked activation were identified. Then, the correlation of the two variables (information transfer or task activation) with memory performance was calculated to determine which was the mediator. Finally, mediation analysis and the Sobel test were performed to judge whether there was a significant mediating effect among information transfer, task activation and memory performance (significance at p < 0.05).

### Diffusion tensor imaging preprocessing

Image preprocessing of diffusion tensor imaging (DTI) data was performed using the PANDA toolbox (http://www.nitrc.org/projects/panda) based on FSL 5.0 for all DTI images (http://fsl.fmrib.ox.ac.uk/fsl/fslwiki) and the Diffusion Toolkit (http://www.trackvis.org/dtk/), including motion and eddy current corrections and coregistration of fractional anisotropy (FA) images to T1-weighted images by affine transformation. The Glasser atlas in standard space was then used to inversely warp images back to individual native space by applying this inverse transformation. This parcellation divided the cortical surface into 360 regions. A streamline was terminated when it reached a voxel with an FA value lower than 0.1 (reflecting low levels of preferred diffusion, often gray matter voxels), when the streamline exceeded the brain mask (i.e., gray and white matter voxels), or when the trajectory of the streamline made a turn sharper than 45 degrees. Streamlines longer than 15 mm were considered in further analyses.

### Structural connectivity definition

Network nodes i and j were defined as being structurally connected when a set of fibers from the total collection of reconstructed streamlines was found that connected them. The weights of the connections were defined as the number of fibers (FN) between two regions. The size of the structurally connected network was 360×360, the same as the size of the information transfer network.

### Analysis of influence of structural connectivities in information transfer

Values of FN>0, FN > 1, FN > 2, and FN > 3 were selected to determine whether SC clearly existed. In the information transfer network of encoding and retrieval, the transfers were divided into direct transfers and indirect transfers according to whether there were SCs between regions. The average value of these two types of transfers and their correlation with memory performance were calculated.

### Validation in psychiatric disorder

The fMRI data of patients with BD were used to validate the results of information transfer in episodic memory. Forty-five BD patients from the OpenfMRI database served as the validation group. The acquisition of fMRI scans and T1-weighted images, the preprocessing of resting-state fMRI and task-state fMRI, and the estimation of FC and task-evoked activation are consistent with the methods described above. Information transfer mapping was used to estimate the region-to-region information transfer in BD patients during encoding and retrieval. Then, the information transfer intensity and traditional attributes (task-evoked activation) were compared between groups.

### Validation through brain state changes

The index obtained by subtracting the BOLD signals at the beginning of each trial from the BOLD signals at the end of each trial could reflect the changes in brain state during the task. The averaged state change was calculated for each type of trial (encoding included “task”, and retrieval included “correct” and “incorrect” outcomes). Then, the correlations between information transfer intensity and brain state changes were analyzed in each subject. It is worth noting that the calculation of the information transfer intensity here involved all of both effective and ineffective transfers. State changes in the brain were determined by the interaction between all regions.

### Validation through artificial neural networks

A three-layer ANN was used to simulate the working mechanism of the brain. The input was the averaged resting-state BOLD signal in each brain region, and the output was the memory score. The first and third fully connected layers were used to simulate the encoding and decoding of signals, and their sizes were 360×360 and 360×1, respectively. The second fully connected layer simulated information transfer between regions, with a size of 360×360. The weights of the first and third layers were trained by the optimizer. The weight of the second layer was determined by our information transfer matrix, which was determined at the beginning and was not included in the training. The type of optimizer was stochastic gradient descent (SGD), and the training ran 1000 times. After the training was complete, effective transfers and an equal number of ineffective transfers (repeat 100 times) were retained in the second layer. Then, we compared the similarity between the output and the original results.

### Data availability

The raw fMRI and MRI data are publicly available at https://legacy.openfmri.org/dataset/ds000030/.

The information transfer mapping code can be found at https://github.com/ColeLab/informationtransfermapping, and the connectome-based predictive modeling code can be found at https://bioimagesuiteweb.github.io/webapp/. Additional post hoc analyses are available from the authors upon request.

## Acknowledgements

This work was supported by the National Key R&D Program of China (grant number 2018YFC0115400), the National Natural Science Foundation of China (grant numbers 81671776, 61727807, 61633018), and the Beijing Municipal Science & Technology Commission (grant numbers Z181100003118007, Z191100010618004).

## Author contributions

Tianyi Yan, Gongshu Wang, Shintaro Funahashi, Duanduan Chen, Bin Wang. and Jinglong Wu designed research; Gongshu Wang, Li Wang and BinWang performed research; Gongshu Wang, Tiantian Liu and Bin Wang analyzed data; Gongshu Wang, Dingjie Suo and Duanduan Chen contributed new analytical tools .Tianyi Yan, Gongshu Wang, Li.Wang. and Bin Wang wrote the paper, Ting Li and Luyao Wang revised the paper.

## Figure Legends

**Figure 2—Supplementary figure 1.**
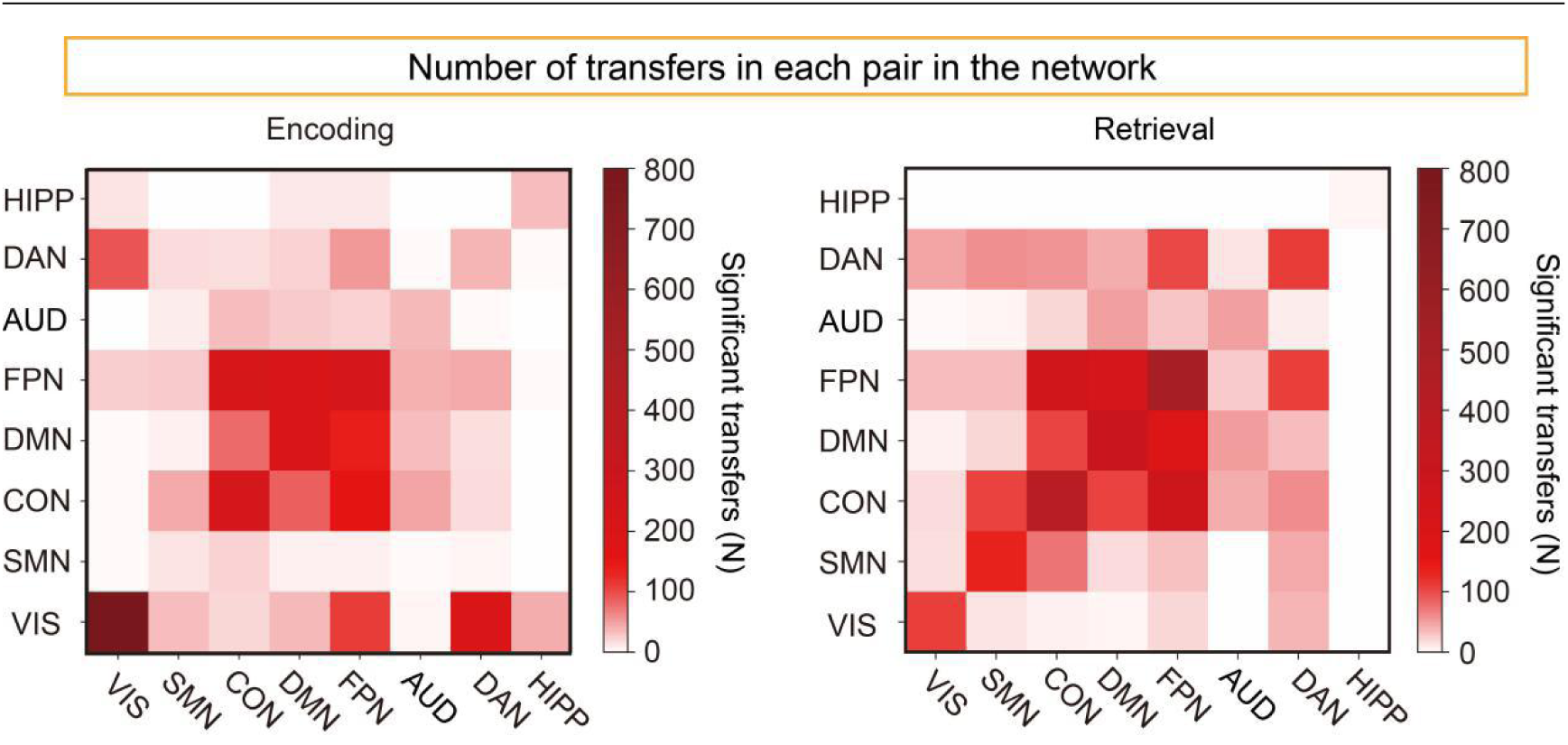
Number of statistically significant region-to-region information transfers by network affiliation during encoding and retrieval. Only the results where the number of transfers of network affiliation was larger than 2 are displayed.

**Figure 5—Supplementary figure 1.**
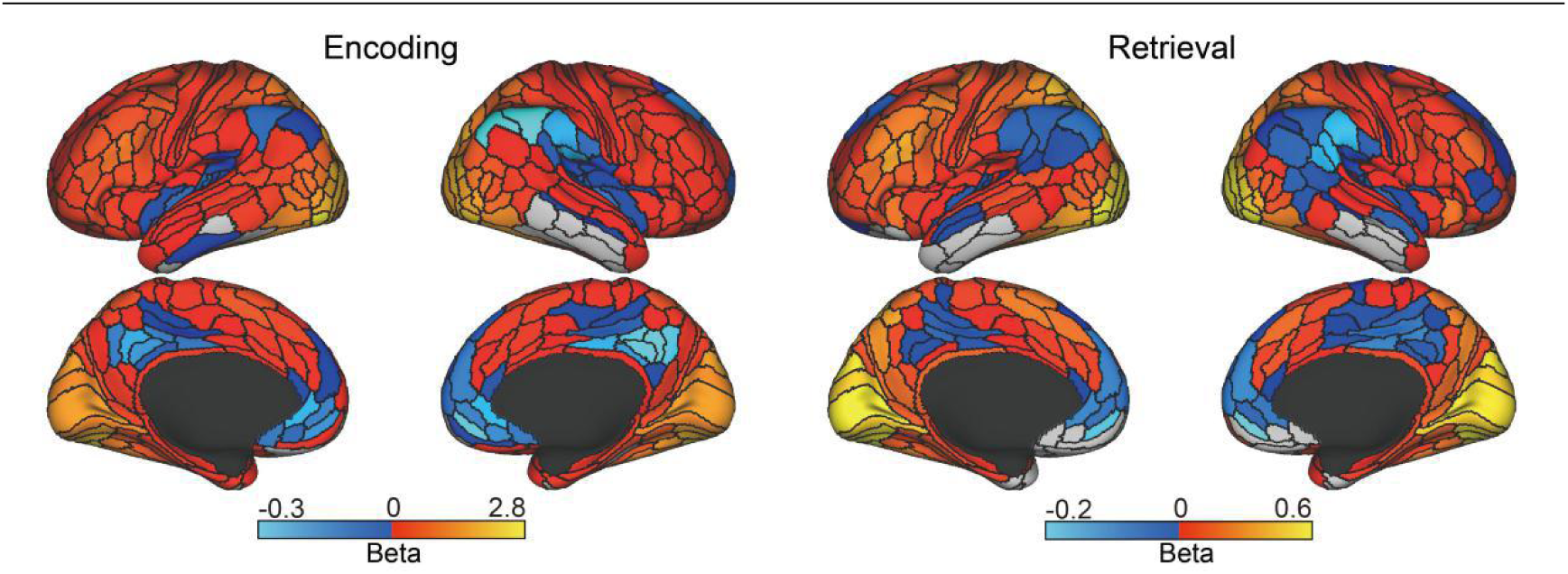
The average beta value of each region during encoding and retrieval processes.

**Figure 5—Supplementary figure. 2.**
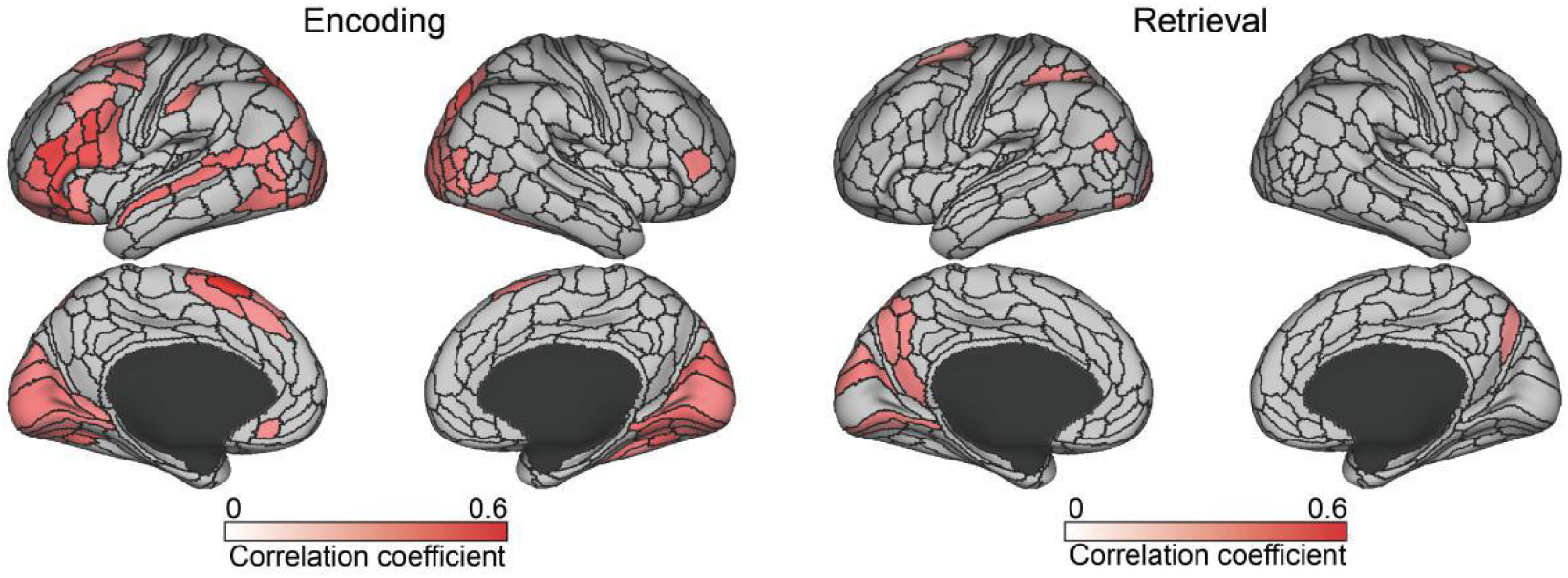
Correlation between task-evoked activation and memory performance in each region. All reported regions were statistically significant at p < 0.01. There were 77 brain regions during encoding and 19 brain regions during retrieval.

**Figure 6—Supplementary figure 1.**
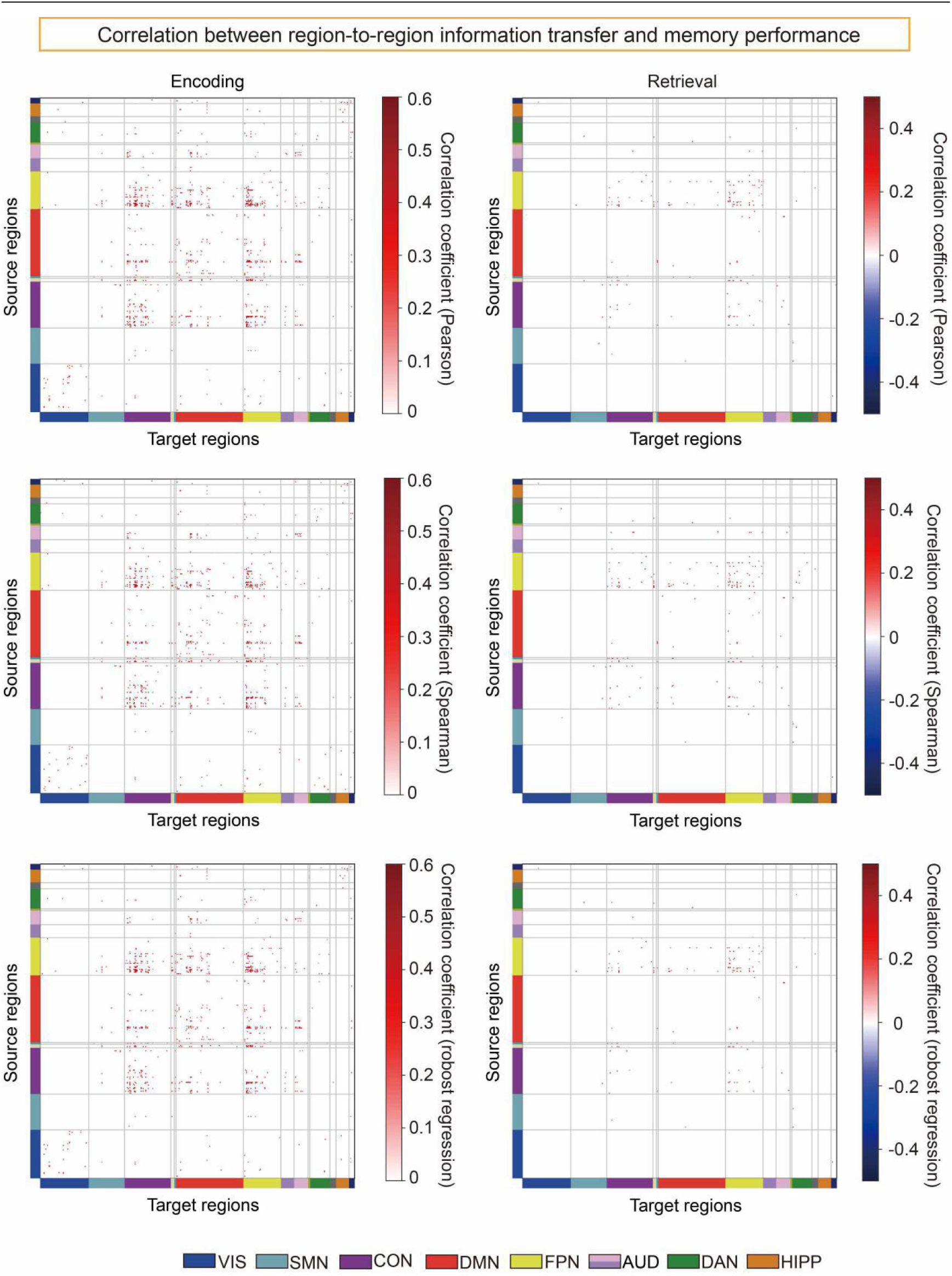
The correlation between each paired information transfer and memory performance during encoding and retrieval processes (p<0.01). Three correlation analysis algorithms were used: Pearson correlation analysis, Spearman correlation analysis and robust regression.

**Figure 7—Supplementary figure 1.**
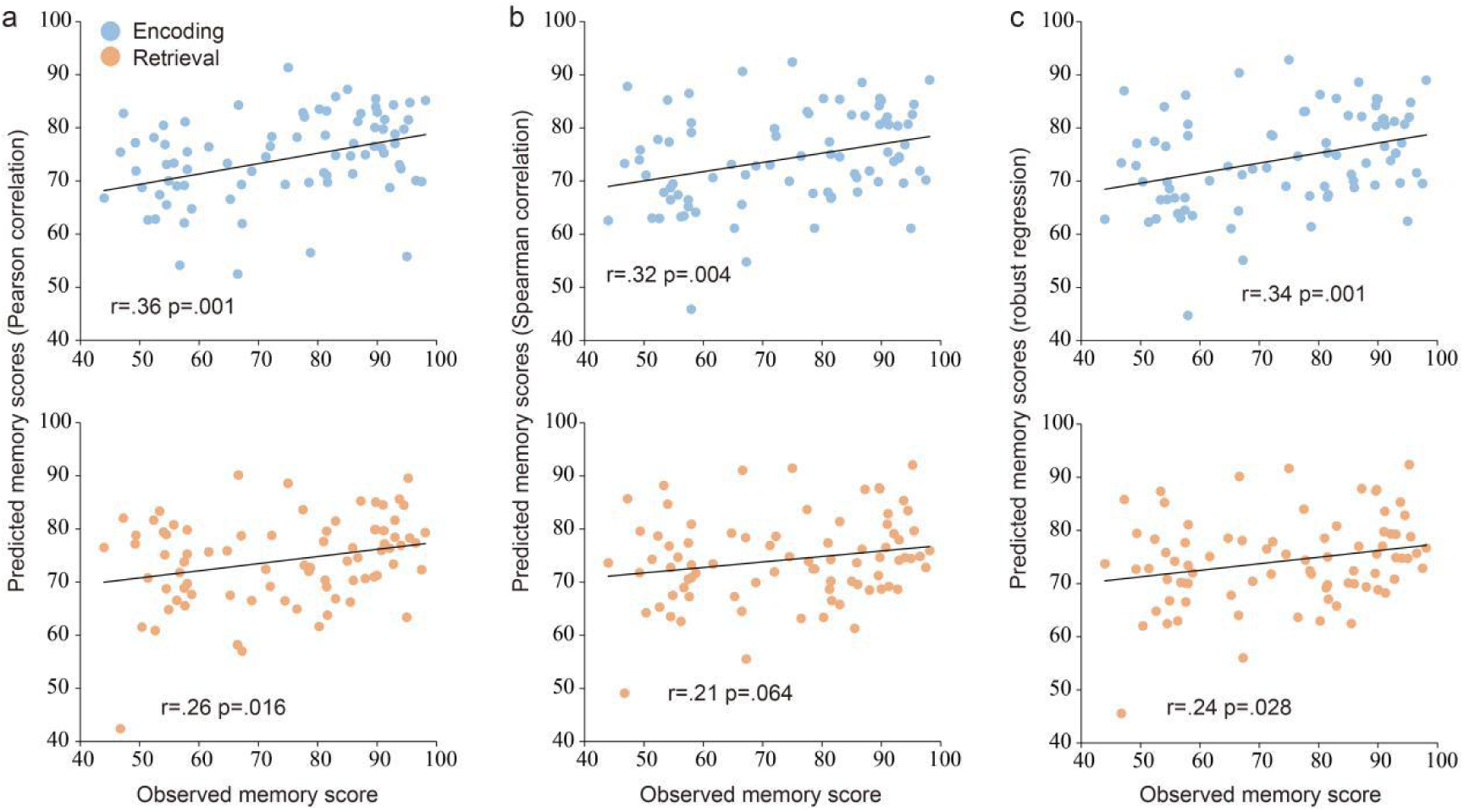
Correlation between predicted and observed memory scores. A tasking-state FC network was employed to predict episodic memory performance. a, b, and c represent three different feature selection methods: Pearson correlation, Spearman correlation, and robust regression (edge-selection threshold of p < 0.01).

**Figure 7—Supplementary figure 2.**
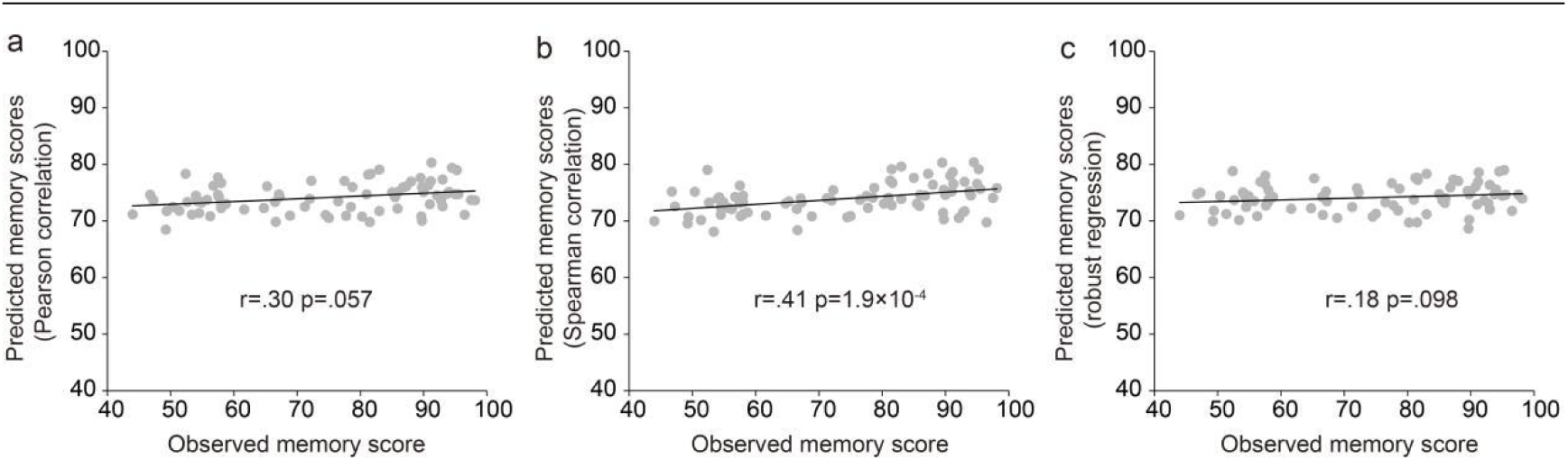
Correlation between predicted and observed memory scores. A resting-state FC network was employed to predict episodic memory performance. a, b, and c represent three different feature selection methods: Pearson correlation, Spearman correlation, and robust regression (edge-selection threshold of p < 0.01).

**Figure 10—Supplementary figure 1.**
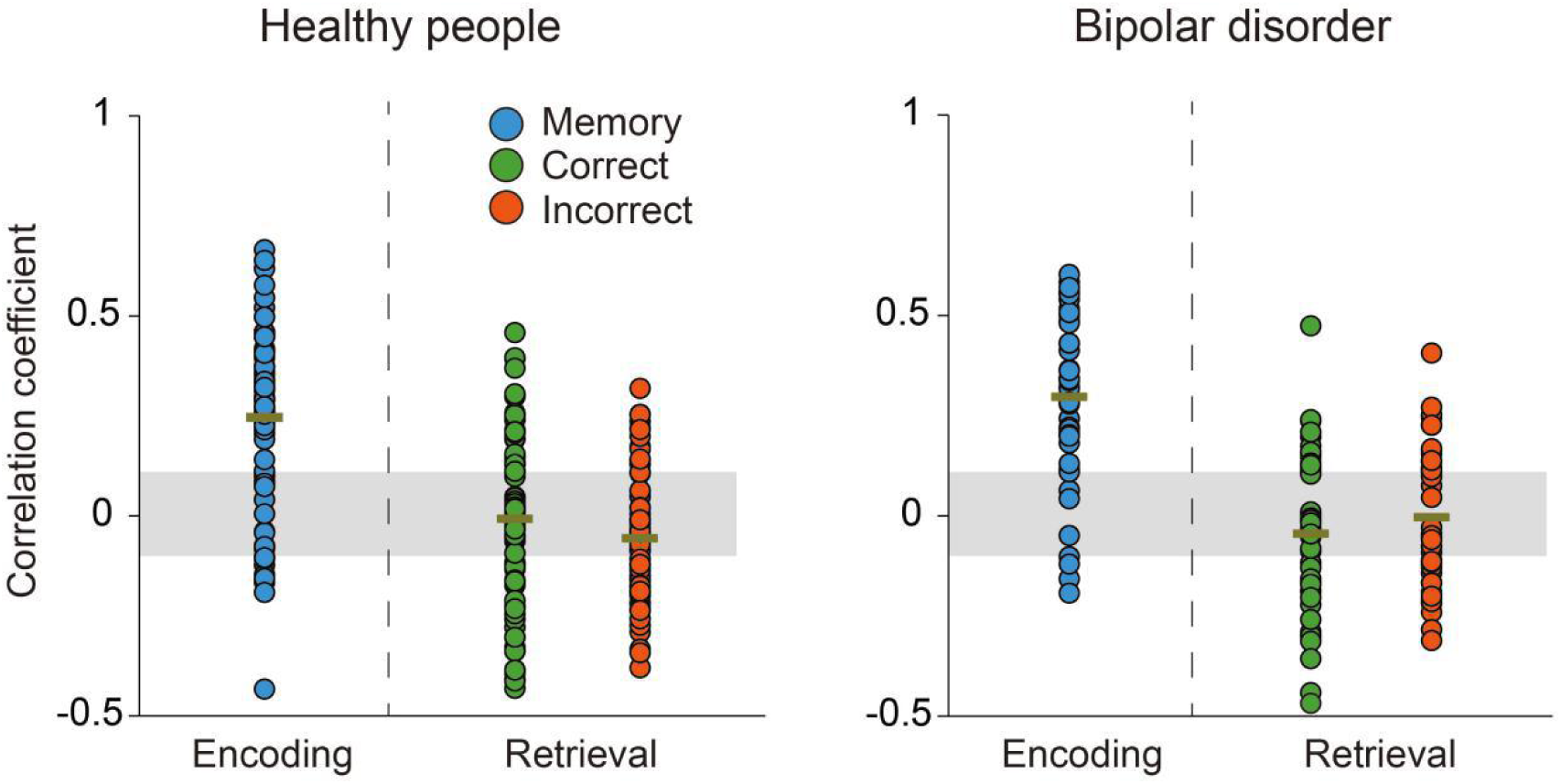
Correlation between regional information transfer intensity and regional activity changes. The gray area indicates that the correlation is not significant (p>0.05).

**Figure 10—Supplementary figure 2.**
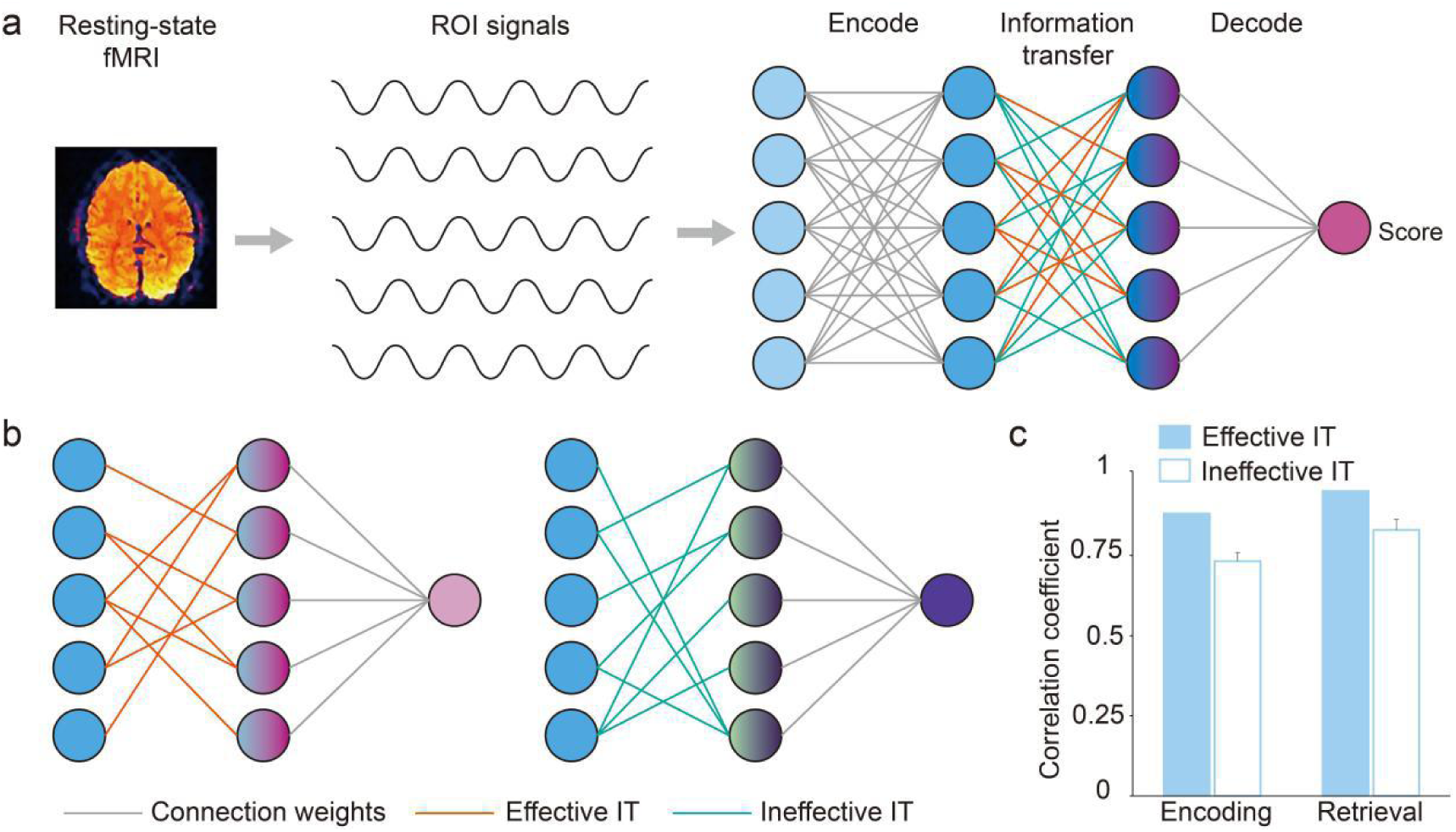
An artificial neural network was used to simulate the functional mechanism of the brain to explore the role of information transfer. **a** The structure of artificial neural network. **b** Effective and ineffective ITs were retained respectively. **c** The influence of effective and ineffective transfers on the model. ROI = region of interest, IT = information transfer

## Source Data

**Figure 2-Source Data File**: T-values of edges in information transfer network(single sample t-test). Values are corrected for a false discovery rate (FDR) of p < 0.05. Information transfer networks estimated by information transfer mapping which is based on activity flow mapping.

**Figure 3-Source Data File**: The regional information transfer percentages were obtained by taking the number of effective transfers (Figure 2) from and to a region and dividing it by the total number of possible transfers.

**Figure 4-Source Data File:** Regions where the information transfer intensity is significantly different between encoding and retrieval (paired sample t-test, p<0.05 with Bonferroni correction).

**Figure 5-Source Data File:** Regions where the information transfer intensity are significantly correlated with task-evoked activation (Pearson correlation analysis, p<0.01).

**Figure 5—Supplementary figure 2-Source Data File**: Regions where the task-evoked activation are significantly correlated with memory score (Pearson correlation analysis, p<0.01). There were 77 brain regions during coding and 19 brain regions during retrieval.

**Figure 6-Source Data File**: Regions where the information transfer intensity are significantly correlated with memory score (Pearson correlation analysis, p<0.01).

**Figure 6—Supplementary figure 1-Source Data File**: Region-to-region information transfers are significantly correlated with memory scores (Pearson correlation analysis, p<0.01).

**Figure 9-Source Data File**: Average intensity of direct and indirect transfers. The type of transfer is defined based on whether there is a structural connection, depending on the number of fiber bundles, between regions.

**Figure 10-Source Data File1:** Comparison between beta-value of healthy subjects and patients with bipolar disorder in each subnetwork (paired t-test, p-values are corrected by Bonferroni).

**Figure 10-Source Data File2**: Comparison between information transfer intensity of healthy subjects and patients with bipolar disorder in each subnetwork(paired t-test, p-values are corrected by Bonferroni).

